# Identification of a large class of cancer–germline microproteins as a source of immunotherapeutic targets

**DOI:** 10.64898/2026.05.18.725836

**Authors:** Marta E. Camarena, Covadonga Vara, Chris Papadopoulos, José Carlos Montañés, Sara Razquin-Sola, Marie Taillandier-Coindard, HuiSong Pak, Markus Müller, Nafiseh Khelgati, Juan Carlos García-Soriano, Puri Fortes, Michal Bassani-Sternberg, Júlia Perera-Bel, M. Mar Albà

## Abstract

Classical cancer germline-antigens (CGAs) are proteins that are expressed in the male germinal line but not in somatic tissues, and that can also become expressed in tumors. However, the vast majority of testis-specific transcripts are long non-coding RNAs (lncRNAs) rather than protein-coding genes. Since recent studies have shown that many lncRNAs contain non-canonical open reading frames (ncORFs) that are translated into small proteins, or microproteins, there could be a large class of non-canonical cancer-germline antigens (ncCGAs) that remains to be discovered. Here, we integrate ribosome profiling from human testis and cancer cell lines with paired tumor/normal transcriptomes from 917 patients across eight common cancer types to define a comprehensive catalog of ncCGAs. This set comprises 235 ncCGAs encoded by lncRNAs or mRNA untranslated regions (5’UTRs and 3’UTRs), compared to 192 canonical CGAs (cCGAs) with similar expression patterns. We show that ncCGAs are evolutionary young, consistent with recent *de novo* emergence in the rapidly evolving male germline. Moreover, a large fraction is expressed across multiple patients and cancer types, indicating recurrent reactivation mechanisms in tumors. We further find that ncCGAs are frequently located in cancer-amplified regions or associated with MYC or E2F-regulated pathways, which may explain their expression in cancer. Finally, we provide strong evidence that a subset of ncCGAs give rise to potentially immunogenic HLA class I bound peptides. Together, our results describe a previously unexplored class of tumor-restricted antigens with potential applications in cancer immunotherapy.

## Introduction

The translation of non-canonical open reading frames (ncORFs) generates thousands of microproteins or peptideins (Mudge et al. 2022; Chothani et al. 2022; Deutsch et al. 2026), whose functions are only starting to be revealed (Chen et al. 2020; Prensner et al. 2021; Sandmann et al. 2023). The translons encoding these small proteins can be detected with high precision using ribosome profiling techniques or Ribo-Seq (Ingolia et al. 2009), a sequencing technique that overcomes previous challenges in annotating small protein coding sequences.

Using high quality Ribo-Seq data from human liver, brain and testis (Wang et al. 2020a), we have recently shown that, in addition to standard testis-specific protein-coding sequences, there is a large number of ncORFs in testis-specific transcripts that show translation evidence but nevertheless lack a protein-coding status (Kaur et al. 2024). It has been known for decades that certain testis proteins can be reactivated in cancer (De Smet et al. 1999; Simpson et al. 2005). The existence of such cancer-germline antigens (CGAs) highlights the similarities between germ-cell formation and tumor development, as some gametogenic pathways - such as immortalization, invasion, induction of meiosis, or migration - are also exploited by cancer cells (Maxfield et al. 2015; Broeils et al. 2023; Laisné et al. 2024). Given these parallels, it is plausible that a subset of the testis-specific translated ncORFs is also expressed in tumors, conforming a previously undescribed class of non-canonical CGAs (ncCGAs). Supporting this idea, we have recently found that about one quarter of the lncRNAs expressed in human hepatocellular carcinoma samples, but not in healthy somatic tissues, are also expressed in testis (Camarena et al. 2024).

Because testis is an immune-privileged organ, with the blood-testis barrier preventing immune infiltration, CGAs can trigger immune responses when expressed in tumors (Traversari et al. 1992; Davis et al. 2004; Whitehurst 2014). Several of them, such as MAGE-A1 and SSX1, have been employed to develop cancer vaccines or antigen-specific T cell-based treatments (Rosenberg et al. 2004; Lu et al. 2017; Sahin et al. 2020). One of the potential limitations of these therapies is the existence of paralogues that are expressed in healthy tissues, which can potentially result in toxicities in healthy organs (Morgan et al. 2013; Leko and Rosenberg 2020). In the case of microproteins encoded by lncRNAs, such paralogues are not expected, since lncRNAs do not typically belong to gene families (Kutter et al. 2012; Ruiz-Orera et al. 2015), reducing the risks of off-target adverse effects. In general, it has been described that many translated ncORFs have emerged *de novo* within primates (Ruiz-Orera et al. 2015; Sandmann et al. 2023). Therefore, it seems likely that CGAs derived from ncORFs will also have a recent evolutionary origin, which would link young gene age with carcinogenic processes.

Here we leverage ribosome profiling data from human testis and cancer cell lines, together with paired tumor/adjacent transcriptomics data from 917 cancer patients, to obtain a comprehensive set of ncCGAs. We find that ncCGAs are abundant, can be shared across cancer types, and are capable of generating HLA-I-bound peptides, opening new paths for immunotherapy applications.

## Results

### Identification of translated ncORFS encoding 235 novel testis-specific microproteins that are reactivated in cancer

The first goal was to characterize ncCGAs resulting from the translation of small ORFs in RNA regions annotated as non-coding and expressed in testis as well as tumors. For this, we used deep Ribo-Seq data from human testis, brain and liver tissue (Wang et al. 2020a), along with extensive RNA-Seq data from healthy and tumor samples (Figure 1A). We considered two main classes of ncCGAs. The class ‘lncRNA-ncCGA’ comprised microproteins translated from small ORFs in transcripts annotated as long non-coding RNAs (lncRNA), processed pseudogenes as well as non-annotated reconstructed transcripts. The class ‘UTR-ncCGA’ corresponded to microproteins translated from upstream and downstream ORFs (uORFs and dORFs, respectively), including cases overlapping the coding sequence (uoORFs and doORFs).

**Figure 1.**
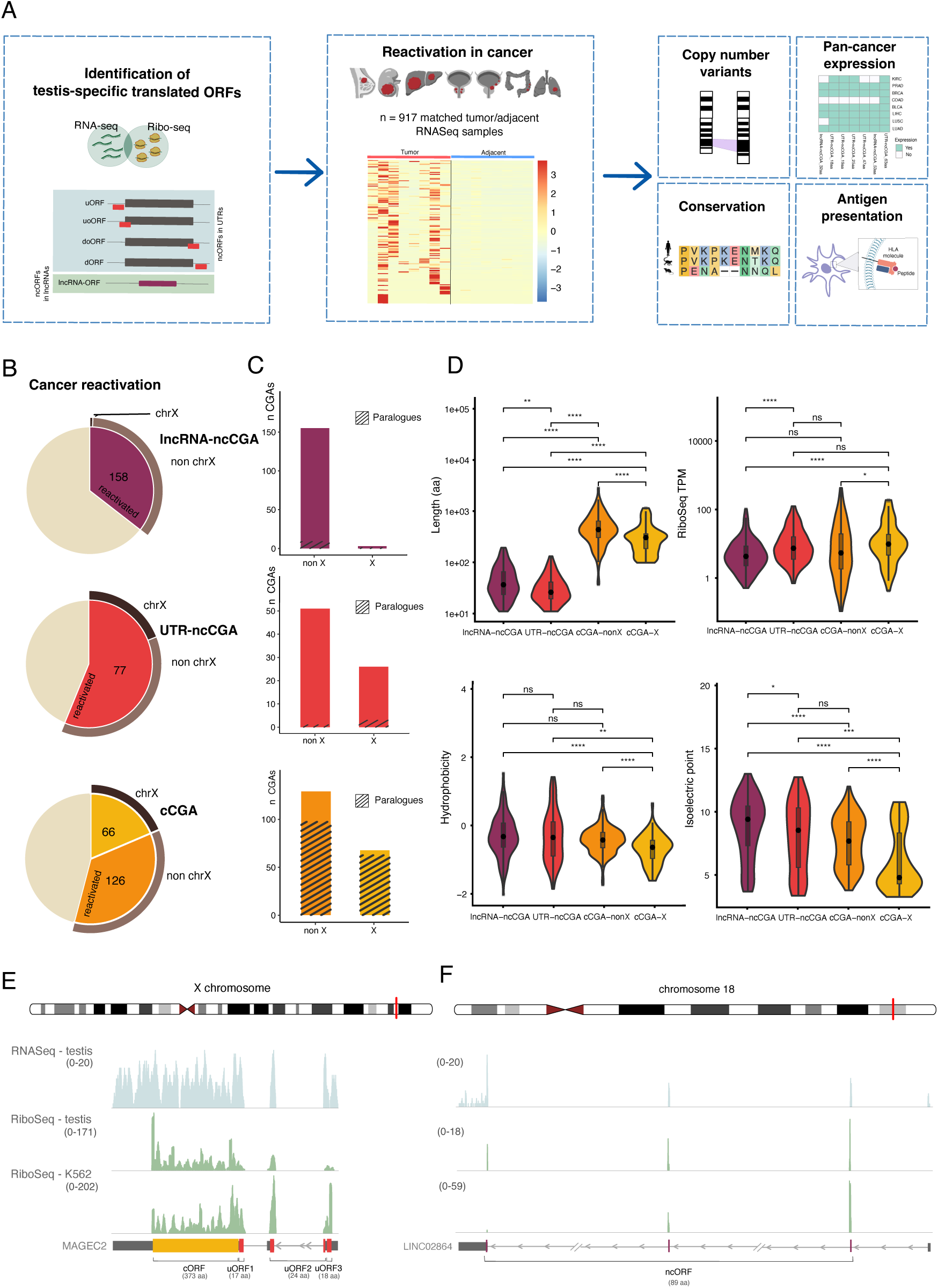
**Reactivation of testis-specific translated ORFs in cancer**. **A.** Overview of the computational workflow used to identify cancer-germline antigens (CGAs) and the main types of analyses. **B.** Proportion of testis-specific translated ORFs reactivated in cancer (CGAs). They have been divided into lncRNA-ncCGAs (highlighted in purple), UTR-ncCGAs (highlighted in red) and canonical CGAs (cCGAs)(cCGA-X highlighted in yellow and cCGA-nonX highlighted in orange). The total number of testis-specific translated ORFs is 445 lncRNA ncORFs, 137 UTR ncORFs and 355 canonical ORFs. The number of ORFs located in transcripts reactivated in cancer is shown in the Figure. **C.** Proportion of CGAs of each type depending on autosome/X localization (non-X and X, respectively) and the existence of paralogues. **D.** Characteristics of cancer-germline antigens (CGA): amino acid length, translation level (Ribo-Seq TPM), isoelectric point and hydrophobicity. **E.** RNA-Seq and Ribo-Seq coverage in human testis and in the K562 cancer cell line of MAGEC2, including the three identified translated uORFs (in red). **F.** RNA-Seq and Ribo-Seq coverage in human testis and in the K562 cancer cell line of LINC02864, with the newly identified translated ncORF (in red).

Translated ORFs were identified with the software RibORF v.2.0 (Ji et al. 2015). We kept ORFs that were exclusively translated in testis, and which were encoded in transcripts lacking expression in somatic tissues from GTEx (median TPM < 0.5 in all somatic tissues analyzed)(Table S1). Reactivation in tumors was assessed by using matched tumor/normal RNA-seq data for a set of 917 patients covering eight cancer types described by The Cancer Genome Atlas Consortium (TCGA) (Weinstein et al. 2013): bladder cancer (BLCA), breast cancer (BRCA), liver hepatocellular carcinoma (LIHC), lung adenocarcinoma (LUAD), lung squamous cell carcinoma (LUSC), kidney renal clear cell carcinoma (KIRC), prostate adenocarcinoma (PRAD) and colon adenocarcinoma (COAD)(Tables S2 and S3). Of the 917 patients, 442 were from TCGA and 475 from 11 additional studies (Yao et al. 2017; Chen et al. 2019; Kannan et al. 2011; Nolan et al. 2022; Wu et al. 2020; Dolgalev et al. 2023; Wang et al. 2020b; Long et al. 2022; Li et al. 2019b; Liu et al. 2016; Repáraz et al. 2022).

Our pipeline resulted in the identification of 235 ncCGAs, comprising 158 lncRNA-ncCGAs and 77 UTR-ncCGAs (Figure 1B). The transcripts encoding them were significantly expressed in one or more patient tumor samples (TPM > 1) and showed null or very reduced expression in the adjacent tissue from the same set of patients (Figures S1 and S2, Table S4). We observed that 27.2% (43) lncRNA-ncCGAs and 62.3% (48) UTR-ncCGAs had start codons other than the canonical ATG (Figure S3). More than half of the cases of UTR-ncCGAs corresponded to uORFs (55%, 42 out of 77 cases)(Table S5); these translated uORFs were located in 31 different cCGA transcripts, including several members of the MAGE family (e.g. MAGEA1 or MAGEC2).

The number of canonical CGAs (cCGAs) identified using the same pipeline was 192, including 53 cancer testis antigens annotated in the CT Database (Almeida et al. 2009). Because there is a large group of cCGAs which is located on the X chromosome (Hofmann et al. 2008; Whitehurst 2014), we separated them into X and nonX cCGAs (66 cCGA-X and 126 cCGA-nonX, respectively) for subsequent analyses. We also checked expression in thymus using RNA-Seq data from 30 different thymus samples. Expression in this organ can induce central tolerance, given its role in eliminating self-reactive T cells (Coulie et al. 2014). Only two cCGAs, GARIN5B and POU4F1, and one lncRNA-ncCGA, MIR924HG, showed significant expression in thymus (47.9, 10.3 and 1.2 median TPM, in the same order)(Table S6).

We quantified the proportion of CGAs that had homologues in the human genome (paralogues) for the different CGA types, as the expression of paralogues has been associated with off-target adverse effects in CGA-based therapies. To account for the fact that microproteins are not annotated, we performed sequence similarity searches against all the microproteins found to be translated in human testis, brain or liver. Unlike cCGAs, the vast majority of lncRNA-ncCGAs and UTR-ncCGAs did not have homology hits to other human proteins or microproteins. This confirmed their sequence uniqueness and potential suitability for the development of safe immunotherapies (Figure 1C, Tables S7 and S8).

### Microprotein CGAs tend to be hydrophobic and/or positively charged

The vast majority of the ncCGAs were smaller than 100 amino acids. In the case of lncRNA-ncCGAs, the median was 37 amino acids and, for UTR-ncCGAs, 26 amino acids (Figure 1D, Table S9). This is in line with previous studies on translated ncORFs (Couso and Patraquim 2017; Ruiz-Orera and Albà 2019; Sandmann et al. 2023). For cCGAs, the size of the proteins encoded in the X chromosome tended to be smaller than those encoded by autosomes (median 305 *versus* 436 amino acids).

We used the mapped Ribo-Seq reads to calculate the translation levels of the different CGA classes (normalized to transcripts per million or TPM). We found that lncRNA-ncCGAs were translated at similar levels as cCGAs-nonX, whereas, in general, UTR-ncCGAs exhibited higher translation levels, comparable to those of cCGAs-X (Figure 1D, Figure S4). Hydrophobicity and isoelectric point values were significantly higher for ncCGAs than for cCGAs, denoting systematic differences in amino acid composition. In general, the properties of the different types of CGAs were similar to the equivalent testis-specific non-tumor reactivated classes (Figure S5). The only exception was the set of cCGAs-X, which were more acidic than their counterparts. We predicted the subcellular localization of the different types of CGAs using DeepLoc 2.1 (Ødum et al. 2024)(Figure S6). For ncCGAs, there was a clear enrichment in the mitochondria (57 lncRNA-ncCGAs and 34 UTR-ncCGAs), whereas only three cCGAs were predicted to be located in this organelle. In contrast, cCGAs were frequently associated with nuclear localization, especially the ones on the X chromosome.

### The translation of ncORFs resulting in microprotein CGAs can also be detected in cancer cell lines

We assessed whether the translation of CGAs detected in testis was consistently observed in two cancer cell lines, chronic myeloid leukemia-derived K562 and human cervical adenocarcinoma HeLa-S3 cell lines, as well as in the human embryonic kidney HEK293T, using available highly quality Ribo-Seq data (Martinez et al. 2020). We detected translation of 41 CGAs in these samples, including 4 lncRNA-ncCGAs and 4 UTR-ncCGAs (Tables S10 and S11). This level of detection was consistent with the number of transcribed CGA-encoding loci. For example, in K562, we could detect expression of the transcripts encoding 81 of the 427 previously defined CGAs by RNA-Seq (TPM>1), and nearly half of these cases, 37, were detected as significantly translated by the analysis of K562 Ribo-Seq data. Considering that the detection of significant translation requires high sequencing coverage to capture the three-nucleotide periodicity pattern of the sequencing reads, this indicates that indeed ncORFs are translated also in the cancer cell lines.

The translation signals of CGAs detected in testis and the cell lines were also comparable. One example was MAGEC2, a known cCGA expressed across multiple cancer types (Hao et al. 2015). We could detect translation of the main coding sequence, as well as of three uORFs, in testis and in K562 cells (Figure 1E). Another example was the lncRNA LINC02864, which contains a ncORF spanning 3 exons and encoding an 89 amino acid microprotein. Again, a similar translation signature was detected both in testis and in K562 cells (Figure 1F).

### Most microprotein CGA are evolutionary young

Previous studies have shown that many translated ncORFs are evolutionary young (Couso and Patraquim 2017; Ruiz-Orera et al. 2018; Sandmann et al. 2023). To determine whether this was also the case for ncCGAs, we developed a pipeline to identify microprotein homologues in *Macaca mulatta* and *Mus musculus* (Figure 2A). First, we used the genome alignments from Zoonomia (Kirilenko et al. 2023) to identify ORFs that would be syntenic to the CGAs defined in human (Figure 2B). Second, conservation of the ncORF was determined at the level of translation. Specifically, we used the Ribo-Seq data from testis, liver and brain for macaque and mouse (Wang et al. 2020a) to assess translational activity at the equivalent ORFs detected by synteny. Subsequently, we ensured that the sequence similarity was significant using BLASTP (E-value < 10^-4^). After applying all these filters, we found 18 lncRNA-ncCGAs (11.4%) and 15 UTR-ncCGAs (19.5%) conserved in macaque, with a single case being conserved also in mouse (Figure 2C, Tables S12 and S13). For comparison, 86.4% of the cCGAs (166 out of 192) showed conservation in mouse using the same methodology.

**Figure 2.**
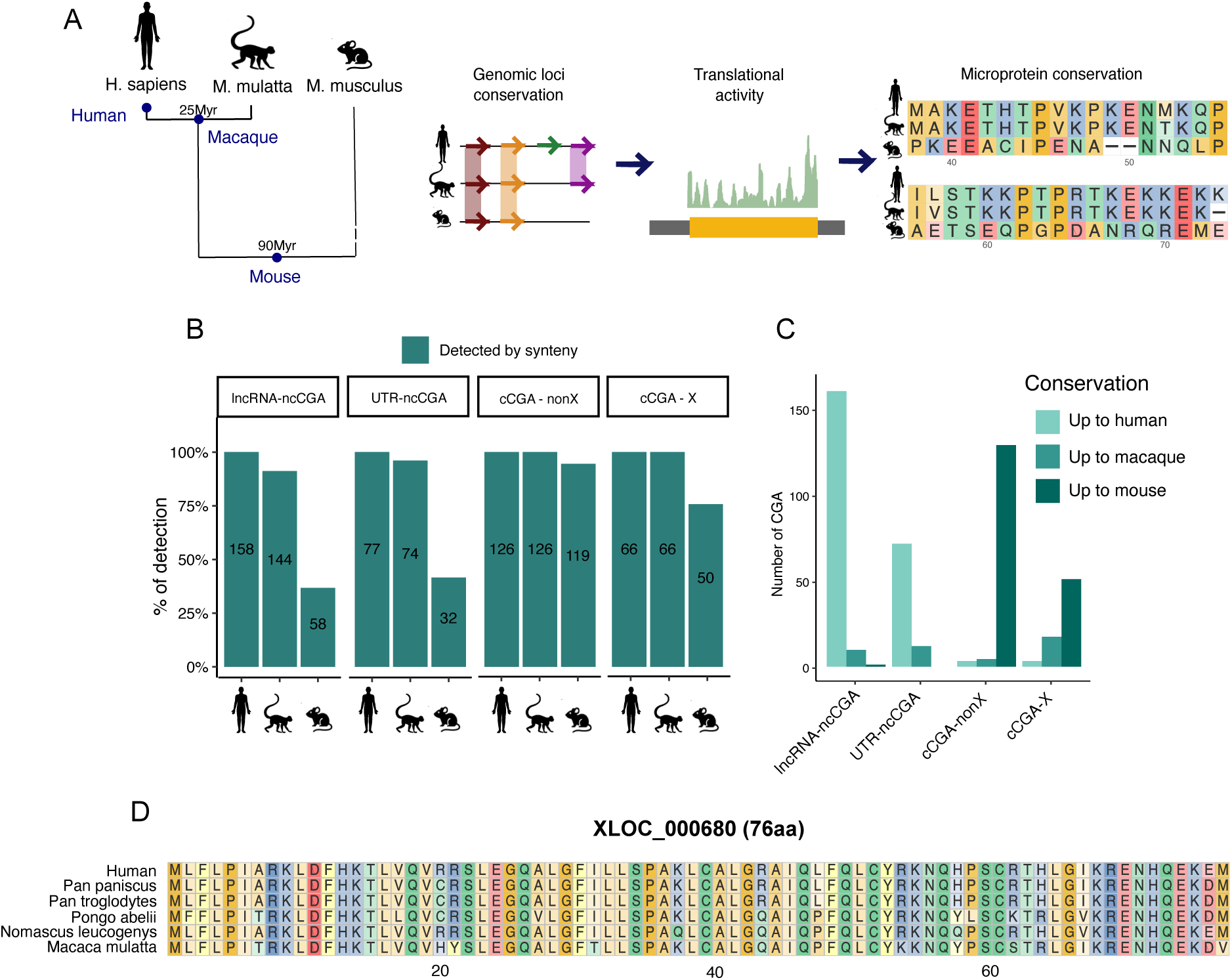
Evolutionary conservation of CGAs in macaque and mouse. **A.** Workflow used to assess evolutionary conservation of CGAs across species. **B.** Proportion of CGAs with conserved ORFs in genomic syntenic regions, stratified by CGA class and species. **C.** Number of CGAs conserved up to human only, up to macaque or up to mouse, using similar Ribo-Seq data for all the species. **D.** Multiple sequences alignment of a microprotein encoded by a newly identified human transcript (XLOC_000680, 76 amino acids).

Therefore, in general, ncCGAs tended to be much younger than cCGAs. The only ncORF conserved in mouse, encoding a 161 amino acid long microprotein, was located in the lncRNA Ensembl gene ENSG00000231110. This analysis also uncovered a novel microprotein of 76 amino acids that is well conserved across primates, and which is encoded by a non-annotated transcript (Figure 2D).

### A large fraction of the ncCGAs is expressed in different cancer types

The number of CGAs detected varied among cancer and CGA types, although liver hepatocellular carcinoma (LIHC) was the one with the highest number of CGAs regardless of CGA type (Figure 3A). Recurrent expression in different cancer types was common. We found that about 45% of the lncRNA-ncCGAs were expressed in more than one cancer type (70 out of 158) and 16 were expressed in at least 4 different cancer types (Figure 3B, Figure S7). For example, LINC02864 was found in all cancer samples except KIRC. The across-cancer distribution of this and other widely distributed cases is shown in Figure 3C.

**Figure 3.**
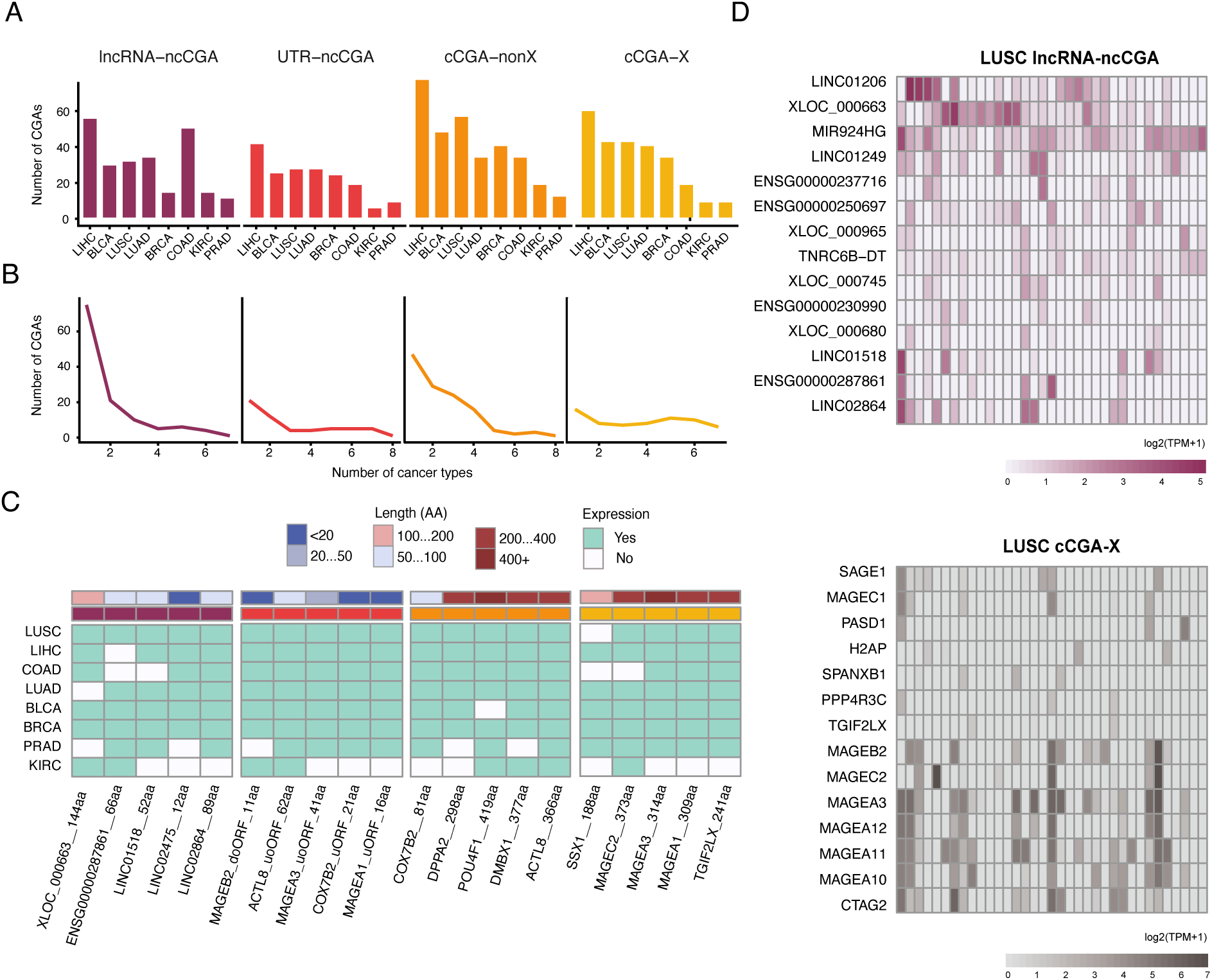
Expression of CGAs across cancer types. A. Number of reactivated CGAs per class and cancer type. KIRC: kidney renal clear cell carcinoma, PRAD: prostate adenocarcinoma, BRCA: breast invasive carcinoma, COAD: colon adenocarcinoma, BLCA: bladder urothelial carcinoma, LIHC: liver hepatocellular carcinoma, LUSC: lung squamous cell carcinoma, LUAD: lung adenocarcinoma. **B. Number of cancer types in which a CGA is expressed**. The data is shown stratified by class. **C. Representative examples of frequently shared CGAs across cancer types.** The CGAs are expressed in 5 or more cancer types. The length of the encoded protein is also indicated. **D. Distribution of commonly found lncRNA-ncCGAs and cCGA-X across patients in LUSC.** The columns represent the different tumor samples; patients are in the same order in the lncRNA-ncCGAs and cCGA-X plots. Color intensity is related to expression level. The CGAs shown are expressed in > 10% of the tumor samples. Transcript names starting with XLOC are non-annotated reconstructed transcripts.

The availability of RNA-Seq data from a large number of patient tumors (917) allowed us to estimate CGA frequencies across patients. We identified 25 lncRNA-ncCGA genes (encoding 39 ncCGA microproteins) and 22 UTR-ncCGA genes (20 ncCGA microproteins) that were expressed in more than 10% of the patients in at least one cancer type (Tables S14 and S15, Figure S8). These highly patient-shared ncCGAs were also frequently expressed in different cancer types (Figure S9). Further, we observed that different lncRNA-ncCGA genes were typically expressed in a different subset of patients, showing limited expression correlation among them in most cancer types (Figure 3D, Figure S10). This mirrors the high tumor heterogeneity observed for other cancer features such as the set of mutated oncogenes/tumor-suppressor genes. In contrast, the expression of cCGA-X genes (such as MAGEs, SAGE1, CTAG2) tended to cluster in the same set of patients, indicating that they are usually activated together.

### ncCGAs are recurrently amplified in cancer

Increased expression of oncogenes in cancer is often achieved through amplification of certain genomic segments (Beroukhim et al. 2010; Zack et al. 2013). We hypothesized that the observed reactivation of testis-specific proteins in tumors could be due, at least in part, to such genome amplifications. To test this, we obtained Copy Number Variation (CNV) data for all eight analyzed cancer types with the program TCGABiolinks (Colaprico et al. 2016). To identify recurrently amplified candidates, we selected genes falling within the top 5% of the segment mean distribution for each cancer type, this cut-off was determined by subsampling random sets of genes.

Using this method, we detected 82 recurrently amplified known oncogenes (25.4% of the analyzed ones), including many previously described cases such as MYC, PIK3CA and SOX2 (Wong et al. 2003; Beroukhim et al. 2010; Rendo et al. 2025)(Tables S16 and S17). Additionally, 30 lnRNA-ncCGA genes (24.6%) and 33 cCGA-nonX genes (26.2%) were also significantly amplified in one or more cancer types (Figure 4A, Figure S11, Table S17). The amplifications were more abundant in LUSC (50 genes), PRAD (47 genes), BLCA (46 genes), COAD (42 genes) and KIRC (42 genes) than in other types of cancer (Figure 4B). The genes detected as amplified overlapped more often than expected with regions described in the Broad TCGA Copy Number Portal (Fisher exact test p-value=0.002), further supporting our findings. One example of a highly amplified lncRNA-ncCGA gene was LINC01206, located in chromosome 3 next to SOX2. This lncRNA was reactivated in LUSC and BLCA, the same cancer types in which we detected the gene amplification. Another case was LINC01344, located in chromosome 1 and significantly amplified in BRCA.

**Figure 4.**
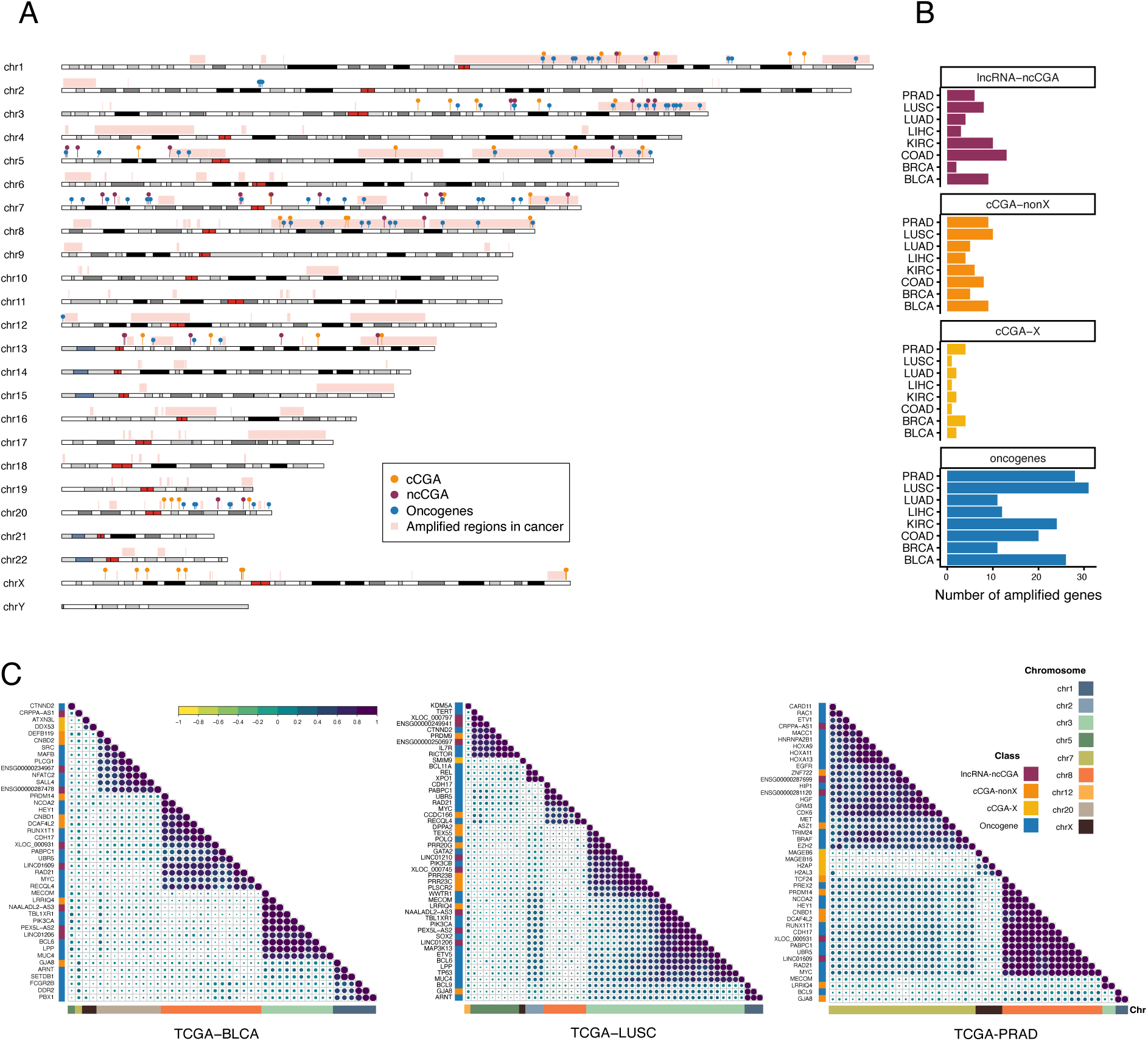
ncCGAs are frequently amplified in cancer. A. Cancer amplified regions and locations of CGAs. Amplified regions from the Broad TCGA Copy Number Portal are highlighted in red. The plot also shows the positions of the genes that were classified as amplified using the program TCGABiolinks. cCGAs are shown in yellow/orange, ncCGAs in purple, COSMIC-defined oncogenes in blue**. B. Number of significantly amplified CGAs by class and cancer type.** The data is shown by cancer type. **C. Level of correlation of amplification status across cancer samples.** Correlation of segment means from TCGA data of genes significantly amplified regions is shown in a scale from-1 to 1, with darker colors for higher values. BLCA, LUSC and PRAD cancer types is shown. Y-axis indicates CGA class, X-axis indicates chromosome.

When we examined the correlation between the amplification values across samples, using the copy number data from the Broad TCGA Copy Number Portal for each locus, blocks of co-amplified genes became evident (Figure 4C, Figure S12). Some of these blocks included both known oncogenes and CGAs of different classes. For example, in chromosome 3, we could distinguish between two groups of amplified oncogenes in LUSC: the first one contained GATA2 and PIK3CB, and the second one PIK3CA, SOX2 and BCL6. The lncRNA-ncCGA LINC01210 was amplified as part of the first group, whereas LINC01206 was present in the second one. Another group was located in chromosome 8 in BLCA and it included MYC, RAD21 as well as LINC01609.

### ncCGAs are associated with cancer-related pathways

To investigate the potential functional roles of lncRNA-ncCGAs, we performed a functional enrichment analysis based on the co-expression profiles of genes across the 917 tumor samples. Focusing on lncRNA-ncCGA genes that were expressed in > 5% of the tumors, we identified 21 out of 50 hallmark gene sets from the Molecular Signatures Database (MSigDB)(Liberzon et al. 2015) that were significantly associated with the lncRNA-CGAs (adjusted p-value < 0.05). The majority of these hallmarks were related to cancer, such as MYC activation, cell cycle targets of E2F transcription factors and progression through the cell division cycle (Figure 5)(Table S18).

**Figure 5.**
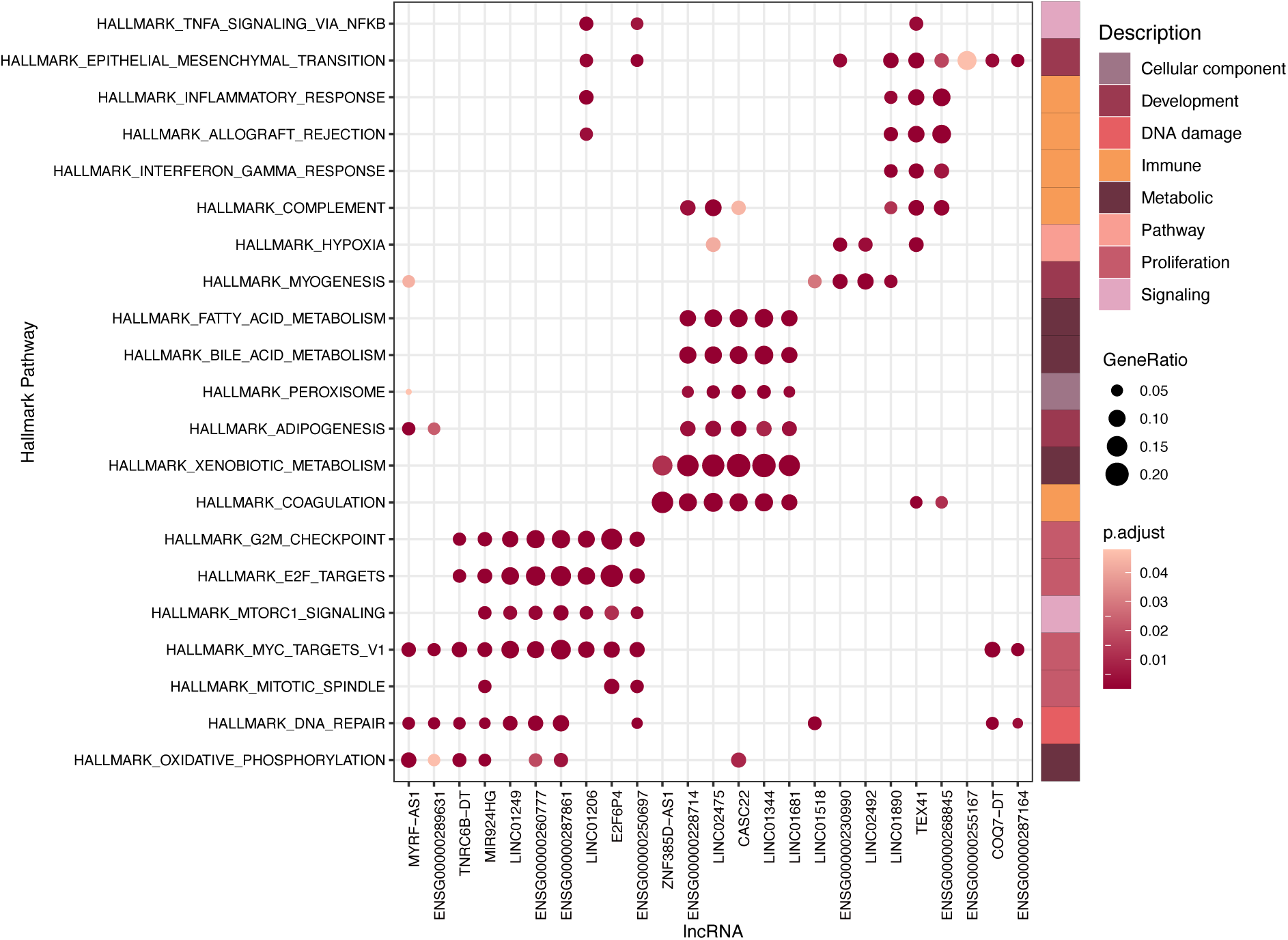
LncRNA-ncCGA functional associations by co-expression with annotated genes. The results were generated by the enricher and clusterProfiler R packages. Only pathways with an adjusted p-value < 0.05 and including at least three lncRNA encoding ncCGAs with a gene ration > 0.05 have been considered. In the Figure, p.adjust refers to the adjusted p-value. GeneRatio refers to the proportion of genes of interest in the gene set out of the total genes of interest.

The group of 12 lncRNA-ncCGA genes potentially activated by MYC included cases found in a large fraction of the 181 LIHC tumors analyzed, including MYRF-AS1 (63 patients), ENSG00000287164 (28 patients) and MIR924HG (22 patients). Only two cases in this cluster, LINC01206 and ENSG00000250697 (expressed in 10 and 7 LUSC samples, respectively), were also located in amplified regions, and so the two activation mechanisms could be largely independent. Additional significant associations corresponded to immune system functions, and to metabolic pathways related to the processing of drugs and lipid metabolism. One example of the latter was LINC02475, a lncRNA-ncCGA expressed in 17% of the hepatocellular carcinoma tumors analyzed. For comparison, we performed a similar functional enrichment analysis for cCGAs. We obtained a similar set of enriched pathways, emphasizing the functional similarities between the two classes of CGAs (Figure S13)(Table S19).

### ncCGAs generate HLA-I-bound peptides

To obtain experimental evidence that ncCGA-derived peptides can bind HLA-I, we examined existing immunopeptidomics data from diverse cancer types, including melanoma, colorectal cancer, breast cancer and lung cancer (9 datasets comprising 266 mass spec files (Tables S20 and S21)(Bassani-Sternberg et al. 2015, 2016; Chong et al. 2018; Gfeller et al. 2018; Chong et al. 2020; Solleder et al. 2020; 2019; Kraemer et al. 2023; Huber et al. 2025), using FragPipe (Kong et al. 2017; Yang et al. 2023; Kall et al. 2007; da Veiga Leprevost et al. 2020). As a database for the searches, we used a human reference proteome concatenated with the 235 ncCGAs detected by Ribo-Seq. The peptide length distribution, and their predicted binding affinities to the respective HLA allotypes, were similar for peptides derived from protein-coding genes in general (PCGs), cCGAs and ncCGAS (Figures 6A and 6B). This analysis recovered 416 canonical cCGA-derived peptides, some of which had been previously reported, including GVYDGEEHSV from MAGEB2 (de Beijer et al. 2022), ESIKKKVL from MAGEC2 (Godelaine et al. 2007), and ALREEEEGV and KVLEYVIKV from MAGEA1 (Ouspenskaia et al. 2021; de Beijer et al. 2022). In addition, we identified 39 ncCGAs-derived peptides (Figure 6C, Table S22). Subsequently, we selected 24 peptides predicted to be HLA-I binders for their respective sample HLA-I alleles (rank<2%), or which had 2 or more identified peptide-spectrum matches (PSM), for further validation (Figure 6D, Table S23). The remaining 15 peptides were not pursued further because they were supported by only a single PSM and were not predicted to bind the corresponding HLA-I alleles, making them more likely to represent false-positive identifications.

**Figure 6.**
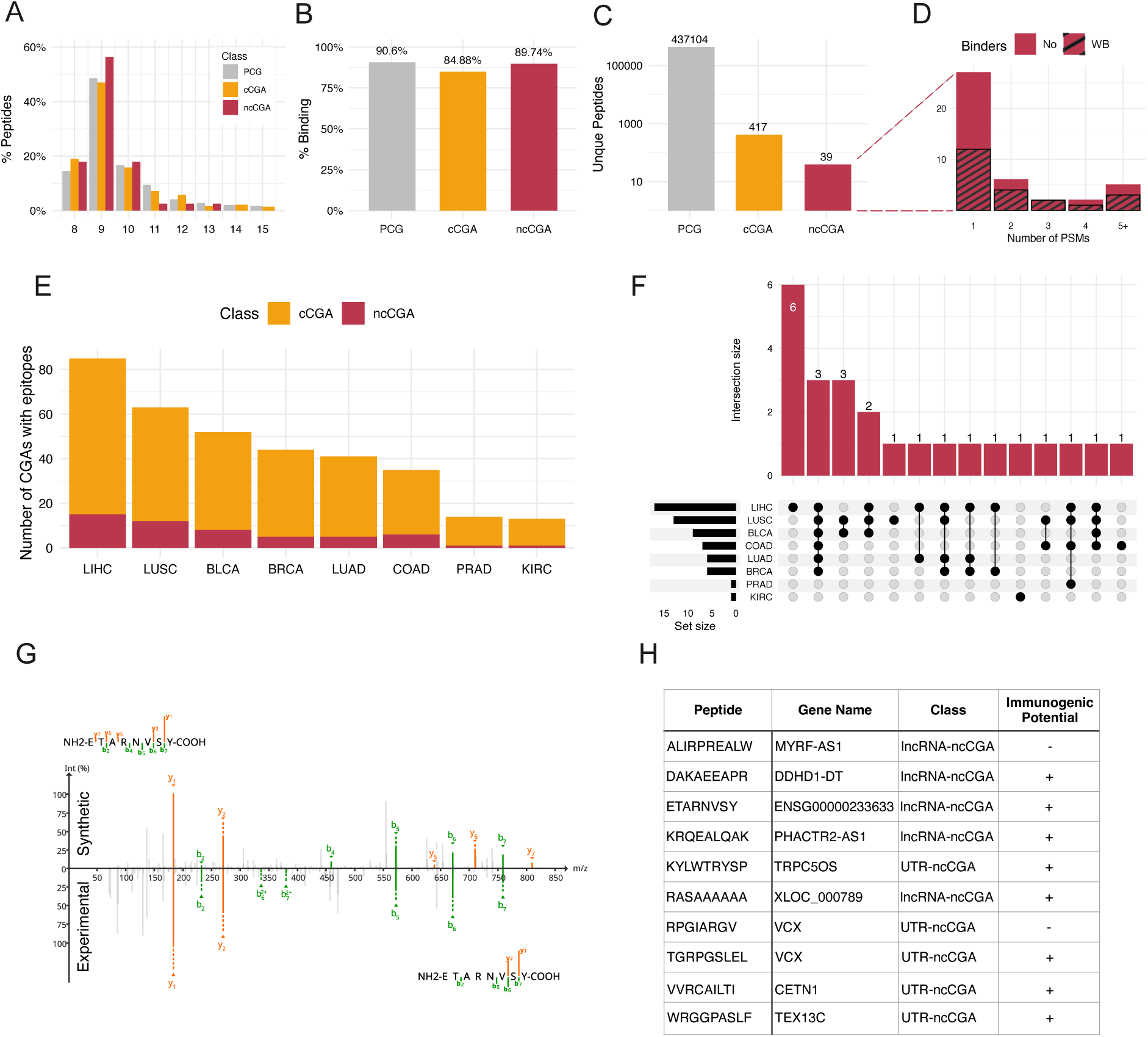
ncCGAs generate tumor-specific HLA-I-bound peptides. A. Peptide length distribution. Length distribution of the peptides detected by immunopeptidomics, separated by class. PCG refers to protein-coding genes from the uniprot database; cCGA to canonical CGAs (also contained in PCGs); ncCGAs to non-canonical CGAs (lncRNA-ncCGAs and UTR-ncCGAs). **B. Binding affinity of peptides detected.** Predicted binding affinity with MixMHCpred for the peptides detected between 8 and 14 Aa long. **C. Number of detected peptides per class. D. Number of peptide-spectrum matches (PSM) and binders per non-canonical peptide.** Number of PSMs and predicted weak binders (rank < 2% from MixMHCpred, labelled as WB) for the 39 non-canonical peptides identified. **E. Distribution of cCGAs and ncCGAs with peptides across cancer types.** Expression of genes encoding cCGAs and ncCGAs harboring peptides detected by immunopeptidomics across the different cancer types analyzed. The data is for 310 cCGA-derived peptides (99 cCGAs) and for 24 ncCGA-derived peptides (23 ncCGAs). **F. Sharedness of ncCGAs with peptides across cancer types. G. Spectra comparison of synthetic and experimental peptide of the lncRNA-ncCGA ENSG00000233633.** The above spectrum corresponds to the synthetic spectra. The below spectrum corresponds to the experimental spectra. **F. Validated tumor-specific non-canonical peptides.** Table with the 10 tumor-specific validated non-canonical peptides and their predicted immunogenic potential (rank < 2%).

The distribution of the filtered peptide candidates (24 derived from ncCGAs and 310 derived from cCGAs) across the 8 TCGA cancer types revealed that hepatocellular carcinoma (LIHC) exhibits the highest number of CGA-derived epitopes, followed by lung squamous cell carcinoma (LUSC) and urothelial bladder cancer (BLCA) (Figure 6E). Conversely, prostate adenocarcinoma (PRAD) and kidney clear cell carcinoma (KIRC) displayed the fewest. This distribution was consistent with the overall number of detected reactivated CGAs in these different cancer types. Several ncCGA-derived peptides were presented across multiple different cancer types (Figure 6F), highlighting their broad potential applicability in pan-cancer immunotherapeutic strategies.

We then performed validations of the 24 ncCGA peptide candidates using synthetic peptides. While the synthesis of six peptides failed, 10 of the 18 MS/MS comparisons confirmed the authenticity of the peptide identifications (Figure 6G and 6H)(Figure S14 and Table S23). Of the 8 that were not confirmed, QSLRSLIIL from LINC01206 showed excellent spectral similarity but failed retention time validation (1058s discrepancy), whereas the others had low spectral similarity. Additionally, two of peptides that failed synthesis were validated against theoretical spectra, resulting in a total of 12 validated peptides (Figure S15)(Table S24).

Two of the 12 validated peptides, ALYTRLPASK and EVGSSLTHTSW, from lncRNAs ENSG00000287175 and ENSG00000267175, were detected in the HLA Ligand Atlas, a compendium of canonical and non-canonical HLA ligands from healthy tissues (Marcu et al., 2021). These hits were to peripheral blood mononuclear cells (PBMC) samples, and thus these two peptides were not considered tumor-specific. Immunogenicity predictions by PRIME 2.1 (Gfeller et al. 2023) flagged 8 of the 10 remaining validated ncCGA-derived peptides (80%) as TCR-recognizable in the context of their respective sample HLA-I alleles (Figure 6H, Table S25). Notably, four predicted immunogenic epitopes bound the common HLA allele A*02:01, which is prevalent globally (Sanchez-Mazas et al. 2024)(VVRCAILTI from dORF of CETN1, KRQEALQAK from the lncRNA PHACTR2-AS1, ETARNVSY from the lncRNA ENSG00000233633, and ALIRPREALW from the lncRNA MYRF-AS1)(Figure S16). Furthermore, two peptides also targeted A*03:01 (DAKAEEAPR, VVRCAILTI), an allele previously linked to cryptic antigens preference (Laumont et al. 2016; Lozano-Rabella et al. 2023).

## Discussion

This study combined Ribo-Seq data from testis and cancer cell lines, together with transcriptomics data from 917 tumor/normal matched samples, to obtain a compendium of non-canonical cancer-germline antigens (ncCGAs). This led to the detection of 235 novel non-canonical CGAs (ncCGAs), about two thirds of which corresponded to microproteins translated from lncRNAs and the rest to translated ncORFs in UTRs. This number exceeds the number of canonical proteins with the same expression distribution (192 cCGAs), emphasizing the pervasiveness of small translated ORF in cancer. Because of their lack of expression in somatic healthy tissues, these ncCGAs represent putative novel targets for antigen-specific immunotherapies such as monoclonal antibodies, T-cell-based therapies or mRNA/peptide vaccines (Sahin and Türeci 2018; Leko and Rosenberg 2020; Paul et al. 2024).

Numerous studies have shown that a relatively large fraction of the mammalian lncRNAs contain small ORFs that exhibit clear signatures of translation (Ingolia et al. 2014; Ruiz-Orera et al. 2014; Ji et al. 2015; Martinez et al. 2020; Chen et al. 2020). Testis exhibits the highest number of tissue-specific lncRNAs (Derrien et al. 2012), many of which contain translated ncORFs (Kaur et al. 2024). LncRNAs are also very abundant in tumors (Iyer et al. 2015; Unfried et al. 2019), being overrepresented in the tumor-specific transcriptome when compared to genes annotated as protein-coding (Camarena et al. 2024). Our study combined Ribo-Seq and RNA-Seq data from both testis and tumors, which allowed us to identify a unique set of 122 lncRNAs that encode 158 different ncCGAs. These lncRNAs showed virtually no expression in thymus samples, decreasing the chances to be recognized by the immune system as self. Some of the ncCGAs were present in a substantial proportion of the patient samples and could thus have an active role in promoting carcinogenesis. One possible candidate is the 89 amino acid microprotein encoded by LINC02864, expressed in more than 10% of the patients in nearly all cancer types analyzed.

Several of the lncRNA-ncCGA genes identified here had been previously found to have roles in cancer. For example, knocking down LINC01518, associated with DNA repair and myogenesis hallmarks in our analyses, was previously shown to decrease cell proliferation in head and neck squamous cell carcinoma (HNSCC)(Swati et al. 2025). Similarly, silencing lncRNA TEX41, associated with various terms related to immune response and tumorigenesis, was reported to inhibit the proliferation, migration, and invasion capabilities of hepatocellular carcinoma (HCC) cells (Li et al. 2025). Both these lncRNAs have been proposed to function as miRNA sponges, and their expression in cancer might result in changes in the expression levels of certain miRNA regulated genes (Wenbo Yang and Yuyou Chen 2023; Swati et al. 2025). Our findings raise the possibility that the microproteins encoded in these transcript also play roles in cancer progression. Finally, the expression of LINC02475 in hepatocellular carcinoma tumors was previously shown to be associated with decreased survival of patients (Lin et al. 2022), which could mean that the RNA, the encoded microprotein, or both, could have a cancer-promoting role.

In addition to translated ncORFs in lncRNAs, the study disclosed many translated uORFs in cCGAs, such as those in the 5’UTR of the *MAGEC2*, *TPTE*, *SPANXB1* and *DCAF8L2* genes. The possible functions of these uORFs are at present unclear, but their translation levels were in general high and comparable to those of the main coding sequence. Other classes of UTR-ncCGAs, such as dORFs, were less abundant and, in general, translated at lower levels. It cannot be discarded that some of the microproteins encoded by these uORFs play yet uncharacterized roles in promoting cancer, perhaps in conjunction with the main protein product of the same gene. One example would be a uORF located in the ASNSD1 transcript, which encodes a microprotein that promotes medulloblastoma cell survival by interacting with the prefoldin-like chaperone complex (Hofman et al. 2024). The catalog of uORFs discovered here can reveal new cancer players and/or therapeutic targets.

In contrast to most canonical CGAs (cCGAs), the vast majority of ncCGAs did not show homology to other protein products of the human genome, and also displayed poor conservation across species. This likely reflects their recent *de novo* origin from previously non-coding sequences, well-documented for other types of microproteins (Ruiz-Orera et al. 2015, 2018; Sandmann et al. 2023; Broeils et al. 2023). The sequence properties of ncCGAs, including a short size, high hydrophobicity and high isoelectric point, are fully consistent with such a *de novo* origin (Montañés et al. 2023). Thus, most ncCGAs have probably originated in the male germline cells of the human or primate lineages. The testis is a rapidly evolving organ in mammals due to competition for reproductive success. It is characterized by rapid changes in gene expression, amino acid substitutions and the formation of new genes, particularly in late spermatogenic stages (Kaessmann 2010; Murat et al. 2023). In this context, the emergence of new transcripts and proteins that help prevent apoptosis and improve cell survival in spermatocytes might have provided a selective advantage. These same genes, however, could have a cancer-promoting effect if they become activated in other cell types (Broeils et al. 2023). Indeed, it has since long been noted that germ and cancer cells share a number of features, such as immortalization and invasiveness, and recapitulation of parts of the gametogenesis program could contribute to tumor formation and progression (Simpson et al. 2005).

An intriguing question is how genes that are only expressed in germinal cells become reactivated in tumors. Hypomethylation of CpG rich promoters of different CGAs, such as MAGEA1, has been shown to play a role in gene activation (De Smet et al. 1999; Loriot et al. 2006). However, this mechanism seems to mainly affect CGAs on the X chromosome (Van Tongelen et al. 2017). Here we found evidence of two other mechanisms leading to the activation of CGAs in cancer. First, we found that CGAs located in autosomes were frequently found in cancer-associated genome amplifications. One possibility is that the genes are accidentally amplified due to their vicinity to oncogenes (Rendo et al. 2025), although, at least in some cases, the amplification could be due to their own oncogenic activity. One possible example of the latter class is CTCFL, a cCGA located in a cancer-amplified region of chromosome 20, known to promote tumorigenesis by activating the expression of hTERT (Van Tongelen et al. 2017). Second, we found that many CGAs were associated with MYC or E2F-regulated pathways. This included 10 lncRNA-ncCGAs that were not located in genomic amplifications, and which might be transcriptionally activated during cancer.

To test peptide presentation by HLA-I, we used of set of high-quality cancer immunopeptidomes. The searches were restricted to ncCGAs with Ribo-Seq evidence rather than on larger collections of putative ORFs, making them very specific. After validation of the spectra with newly synthesized peptides, we could confirm the presentation by HLA-I of 10 ncCGA-derived peptides. Two of these peptides were part of a microprotein of 31 amino acids translated by an uoORF in the gene VCX. This gene belongs to a family of proteins that share a high degree of sequence identity, including VCX2 and VCX3A. The translated uoORF, however, was not identified in other family members. The VCX gene was expressed in more than 10% of LIHC and LUSC patients, and thus it could be a useful target for these cancer types. Considering that ncCGAs are typically expressed by a subset of the patients, it seems likely that many more ncCGA-derived HLA-I ligands will be discovered in the future when more samples are analyzed.

Our results support the notion that ncORFs contribute in a significant manner to the tumor-specific antigen landscape (Erhard et al. 2018; Laumont et al., 2018; Chong et al. 2020). Additionally, the high hydrophobicity of ncCGAs is expected to increase the likelihood of HLA-I peptide presentation (Huang et al. 2011), and C-terminal hydrophobicity has been linked to increased protein degradation (Kesner et al. 2023). Recent i*n vitro* and *in vivo* studies have demonstrated that tumor-specific peptides encoded by ncORFs can elicit immune responses involving CD8+ T cells (Camarena et al. 2024; Raja et al. 2025; Ely et al. 2025). This has also been observed for classical cancer-germline antigens. In the early nineties, a peptide present in MAGE-A1, as well as MAGE-A3 (which show 73% identity), was shown to be recognized by autologous cytolytic T lymphocyte (CTL) clone (Traversari et al. 1992; Gaugler et al. 1994). Since then, other antigens derived from CGAs have been identified and used to induce an immune response in patients (Davis et al. 2004; Rosenberg et al. 2004; Lu et al. 2017), although in some cases toxicities due to the presence of paralogs expressed in healthy tissues have been described (Morgan et al. 2013; Leko and Rosenberg 2020). More recently, RNA-based vaccines expressing tumor-associate antigens have been developed (Sahin et al. 2020). We found that, unlike cCGAs, ncCGAs tend to have unique sequences, making them potentially safer for the development of immunotherapies.

This study has several limitations. The translation evidence for most ncORFs was obtained from testis Ribo-Seq data, with only a few high-quality available samples to check their translation in cancer. Whereas the results indicate consistency in translation patterns in the two types of cells, more direct evidence will require performing additional Ribo-Seq experiments. The question also remains opens as to how many of the ncCGAs are stable and functional. The pan-cancer analysis helped identify several strong functional candidates, such as those expressed in a large number of patient tumor or associated with oncogenic pathways by co-expression analyses. This suggests that, even if young, some of these proteins might have already transitioned to a functional state, perhaps in the context of the high selective pressures associated with the male germinal line.

In summary, this study provides the first comprehensive description of a large set of testis-specific microproteins that become reactivated in tumors. These ncCGAs are evolutionary young, lack homology to other human gene products, can be shared across patients and cancer types, and generate HLA-I-bound peptides. The work paves the way for the development of immunotherapies based on ncCGAs.

## Material and Methods

### Sequencing datasets

We used public raw Ribo-Seq and matched paired-end stranded RNA-Seq data for three organs (brain, liver and testis) in three mammals (human, macaque, mouse)(Wang et al. 2020a) to predict translated ORFs. They are available from ArrayExpress under the accession number E-MTAB-7247.

For the cancer data, we downloaded matched tumor and adjacent tissue RNA-Seq data from for 917 patients from 12 different sources (Tables S2 and S3). The Cancer Genome Atlas (TCGA) includes data from 8 common cancer types: breast invasive carcinoma (BRCA), liver hepatocellular carcinoma (LIHC), bladder urothelial carcinoma (BLCA), lung adenocarcinoma (LUAD), lung squamous cell carcinoma (LUSC), kidney renal clear cell carcinoma (KIRC), colon adenocarcinoma (COAD), prostate adenocarcinoma (PRAD). Eleven additional paired RNA-Seq datasets for the same cancer types were downloaded from the Gene Expression Omnibus (GEO) and the European Nucleotide Archive (ENA) databases (see Table S2 for references and accession numbers).

Ribo-Seq and RNA-Seq data from human embryonic kidney cell line HEK293T, chronic myeloid leukemia-derived cell line K562 and cervical cancer-derived HeLa-S3 cell line (Martinez et al. 2020) were downloaded from GEO under the accession GSE125218.

### *De novo* transcriptome assembly

For each RNA-Seq sample, we first assessed the quality of the RNA-Seq data with FastQC (v0.11.5) and FastQScreen (v0.14.0) software (Wingett and Andrews 2018), and then aligned the sequencing reads to the reference genome using two-pass alignment with STAR (v2.7.1)(Dobin et al. 2013). The reference genomes used were: GRCh38/p13 for human; GRCm39 for mouse; Mmul_10 for macaca. Only uniquely mapped reads were considered.

We reconstructed novel, non-annotated, transcripts using StringTie (v2.0) in conservative stranded mode (Pertea et al. 2015). Gffcompare was used to merge the novel transcripts from different samples or tissues, based on overlapping genomic coordinates (Pertea and Pertea 2020). We exclusively selected those not overlapping with any annotated transcript using BEDTools (v2.2.1)(Quinlan and Hall 2010). We employed GENCODE human (v47) and mouse (vM32), and the Ensembl transcriptome for macaque (v10.111).

### Human ribosome profiling sequencing data analysis

The ribosome profiling (Ribo-Seq) reads for brain, liver and testis samples were mapped to the customized reference transcriptome. We predicted translated ORFs with RibORF (v2.0) (Ji 2018), which is based on measuring the three nucleotide periodicity and homogeneity of the mapped Ribo-Seq reads. We selected ORFs starting with NTG, longer than 30 nucleotides, with more than 10 reads and a RibORF score > 0.6. For all non-canonical ORFs, the starting amino acid was set as Methionine, independently of the codon, assuming a mismatch of the tRNA-Met (Peabody 1989).

The analysis of ribosome profiling data from cancer and embryonic cell lines was done following the same Ribo-Seq pipeline as for testis samples. Out of the 25 available ribosome profiling samples (12 from HEK293T, 8 from HeLa-S3 and 5 from K562), 14 met our quality criteria and were used for the analyses. Quality control indicated that HEK293T had the highest sequencing depth. However, the K562 cell line, derived from chronic myeloid leukemia, exhibited the greatest number of translated CGAs. Although different treatments were applied across the cell lines, all three K562 samples analyzed were not treated. In total, 33 canonical CGAs and 8 non-canonical CGAs —comprising 4 lncRNA-CGAs and 4 UTR-CGAs—were translated in at least one of the cell lines.

### Testis-specific translated ORFs

After predicting translation from each of the Ribo-Seq samples, we kept ORFs translated in at least one human testis samples but in none of the samples from brain or liver (Figure S17). A minimum of 1 transcript per million (TPM) in one testis sample RNA-Seq data was also required to consider a gene as expressed in testis. A maximum of 0.5 TPM RNA-Seq per brain or liver sample was permitted to classify the gene as testis-specific. Novel transcripts shorter than 300 bp or longer than the longest annotated tumor lncRNA (KCNQ1OT1, 91666 nucleotides) were not considered for further analysis. Expression data from the Genotype-Tissue Expression (GTEx) project (v10)(Consortium 2013) was also collected and used to discard genes expressed in somatic tissues. In addition to the previous criteria, genes with a median expression higher than 0.5 TPM in any non-germinal tissue were discarded. This was applied to the annotated transcripts but not to novel transcripts, as GTEX is not informative in this case. Additionally, we used BLASTP with short mode and an E-value cut-off of 0.0001 (Altschul et al. 1997a) to filter out translated ncORFs with homology to annotated proteins; this might be especially relevant for pseudogenes classified as lncRNAs (Figure S18).

After applying all these filters, we recovered 355 testis-specific translated canonical ORFs (annotated coding sequences), 445 lncRNA ncORFs and 136 UTR ncORFs.

### Tumor/adjacent tissue matched RNA-Seq data

For TCGA data, files containing information on the mapped RNA-Seq reads were downloaded from the Genomic Data Commons Data Portal and transformed to FASTQ format using SamToFastq from Picard Toolkit (v2.25.1). For the rest of GEO and ENA datasets, FASTQ files were directly downloaded. All RNA-Seq reads were quality assessed using both FastQC (v0.11.5) and FastQScreen (v0.14.0) software. All selected samples passed the quality control.

We performed several quality checks of the RNA-Seq matched data. We run a Principal Component Analysis (PCA) with the 100 most variable genes per cancer type to validate that adjacent and tumor samples were properly separated (Figure S19). Additionally, we checked the expression of some tumor biomarkers described to be differentially expressed in tumor and adjacent samples. The selected biomarkers were BRCA1, BRCA2, PTEN for breast cancer (Jin et al. 2022); TERT, THBS4, AFP, MTM1 for hepatocellular carcinoma (Piñero et al. 2020; Cancer Genome Atlas Research Network 2017; Sun et al. 2018; Mao et al. 2012); PLK1, CDK1, CNN1 for bladder cancer (Hussain et al. 2017); ABCA8, ADAMTS8, CEP55, PYCR1 for lung cancer (both adenocarcinoma and squamous cell carcinoma)(Wang et al. 2022); TRIB3, STMN4, FAM135B for colon adenocarcinoma (Wang et al. 2022); AR, GOLM1, PCA3 for prostate adenocarcinoma (Conteduca et al. 2019; Varambally et al. 2008); and CAV1, ALB, VEGFA for kidney cancer (Li et al. 2023). There were clear differences in the PCA and the biomarkers between tumor and adjacent samples, validating the consistency of the tumor/adjacent tissue datasets (Figure S20).

### Identification of cancer-germline antigens (CGAs)

We next determined which of the genes encoding testis-specific ORFs identified earlier were reactivated in tumors using the matched tumor/adjacent tissue RNA-Seq data. To quantify gene expression, we employed an annotation file containing the longest transcript per gene as the representative transcript. This file included both annotated and novel transcripts. We then quantified gene expression in the 917 tumor and adjacent samples using featureCounts (Liao et al. 2014). We used transcripts per million (TPM) as a measure of gene expression in the samples from each of the 8 cancer types (Table S2). Cancer-germline antigens (CGAs) were selected as to have testis-specific expression, as defined earlier, and also be expressed in one or more tumor samples, of one or more cancer types, with TPM > 1.

We also required that the average gene expression in the tumor samples was at least three times larger than the average expression in the corresponding matched adjacent samples in the same cancer type. We employed the following equation:

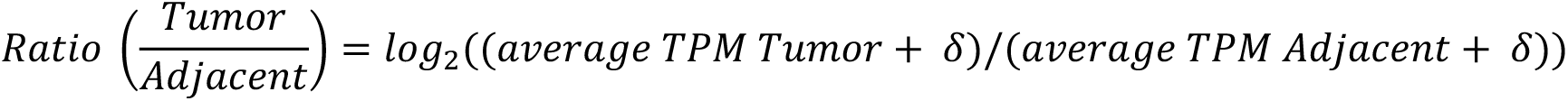

where δ equals 10^-6^

We required that Ratio(Tumor/Adjacent) > 1.585 (Table S4). Although the adjacent tissue might in some cases be contaminated with tumor material, the differences between tumor and adjacent were in general very notable in all selected cases. On average, expression in the tumor samples was 28 times expression in adjacent samples.

The final set of CGAs comprised 192 cCGAs – of which 66 are located in genes in chromosome X and 126 in autosomes-, 158 lncRNA-ncCGAs and 77 UTR-ncCGAs. For each CGA and cancer type, we determined the percentage of patients that expressed the CGA genes in the tumor samples (RNA-Seq TPM > 1) and, from those, the number in which the TPM in the corresponding adjacent sample was smaller than 0.1 (Table S15).

### Identification of CGA paralogues

To determine whether CGAs had paralogues encoded in the human genome, we combined protein sequences from GENCODE version 47 with the translated non-canonical sequences identified in this study from testis, liver, and brain tissues (Wang et al. 2020a). These amino acid sequences were used to construct a database to perform BLAST searches (Altschul et al. 1997). CGA sequences were queried against this database using the blastp-short option (BLAST+ v2.12), with an e-value threshold of 0.0001, to accommodate for the short size of the microproteins. If a CGA sequence showed homology to any protein in the database other than itself, it was classified as having paralogues.

### CGA properties

We extracted the protein sequences of all CGAs and calculated hydrophobicity and isoelectric point, using the hydrophobicity and pI functions of the Peptides R package (Osorio et al. 2015). We also predicted the subcellular localization of the different types of CGAs using DeepLoc 2.1 (Ødum et al. 2024). If several localizations were reported, we chose the one with the highest probability assigned.

We predicted the immunogenicity of the peptides derived from CGA using PRIME2.1 (Gfeller et al. 2023) with a %rank lower or equal to 1 for HLA-A02:01 (Schmidt et al. 2021). We found 117 significant putatively immunogenic non-redundant peptides in 158 lncRNA-CGAs, 36 in 77 UTR-CGAs and 2,032 in 192 cCGAs.

### Thymic expression

We checked the expression of the CGAs in the thymus using 30 publicly available thymic epithelial cells (TEC) samples from two different datasets: GSE127825 (Laumont et al. 2018; Cleyle et al. 2022) and GSE201719 (Carter et al. 2022). We followed the same RNA-seq pipeline as for the other samples here, and we calculated the median expression in thymic cells in TPM units. Genes with a median TPM > 1 in the TEC samples were classified as expressed in thymus.

### Conservation of microproteins in primates and mammals

To assess whether cancer-testis antigens derived from microproteins are conserved in other species, we employed a synteny-based homology detection approach. For the canonical CGAs, we used orthologous gene annotations from the Zoonomia Project for mouse (*Mus musculus*, mm10) and macaque (*Macaca mulatta*, rheMac10), using the human genome (hg38) as the reference (Zoonomia Consortium 2020). For ncORFs, which are not included in Zoonomia annotations, we performed independent orthology inference using TOGA with default parameters and the Zoonomia whole-genome alignments generated with Progressive Cactus (Armstrong et al. 2020). We observed limited detectability of genes located on the X chromosome, which we hypothesized is due to sequence homology among gene family members, which makes the orthology assignment more difficult. To try to identify missed cases, we complemented the TOGA-based approach with pairwise sequence-based orthology detection using proteinOrtho (Lechner et al. 2011). To evaluate whether the short length of ncCGAs limited synteny detection, we stratified them by length and compared detection rates in macaque. We observed similar detection rates for ncCGAs shorter than 20 amino acids (aa)(105 detected and 8 non-detected) than for protein equal or longer than 20 amino acids (382 detected and 23 not detected, Chi-square test, p = 0.74).

Next, we used Ribo-Seq data from three tissues in macaque and mouse (Wang et al. 2020a) to assess translational activity at the genomic loci corresponding to our set of ORFs. Using RibORF, we calculated per-sample offsets and then pooled the data across samples to maximize detection sensitivity. Translated ORFs in the other species were those with longer than 30 nucleotides and with a RibORF score higher than 0.5. We validated the results by performing sequence similarity searches with BLASTP+ (v2.12)(blastp-short option and an E-value threshold of 0.0001). This ensured that the microprotein detected as homologous in the other species had significant sequence similarity.

According to these results, we categorized the 427 CGA in different age groups: 1. Human: no homologues in the other species; 2. Macaque: homologues only in macaque; 3. Mouse: homologues in mouse. The number of CGAs in each class can be found in Table S12. Results per each individual CGA can be found in Table S13.

### Copy Number Variants

We obtained Copy Number Segment data, generated using the DNAcopy workflow, from primary tumor samples for the eight analyzed cancer types (BLCA, BRCA, LIHC, LUAD, LUSC, KIRC, PRAD, and COAD). This data was retrieved using the R package TCGABiolinks (Colaprico et al. 2016). Amplification levels were reported as log2 ratios of copy number for each genomic segment. In addition to the CGAs, we analyzed the data of a set of 232 oncogenes from the COSMIC database (Sondka et al. 2024)(v102, last accessed 26^th^ May 2025)(Table S16). For each gene category, we conducted 500 iterations of random subsampling, retrieving genes from the genome with a length distribution matching that of CGAs. The overlap between the random sets and the CGAs was minimal: 0.02% for lncRNA-ncCGA and 0.87% for CGA-nonX on average. For CGA-X it was 15.7%, due to the requirement of being in the X chromosome. We retrieved as significant those genes falling within the top 5% of the segment mean random distribution for each category. On average across cancer types, 30% of lncRNA-CGA (sd = 22.3), 27% of cCGA-nonX (sd = 21.8), and 12% of cCGA-X genes (sd = 12.6) fell within this top 5% threshold.

To further validate our findings, we retrieved copy number data from the Broad TCGA Copy Number Portal (Tumorscape.org). We used the Tumorscape 1.2.1 dataset (“2015-06-01 stddata 2015_04_02 arm-level peel-off”) for all eight analyzed cancer types. We then assessed whether CGAs identified in our study were located within regions marked as amplified in this reference dataset. As a negative control, we performed 1,000 random iterations to test whether such overlap could occur by chance.

Using Fisher’s exact test on the resulting contingency tables, we found that CGAs overlapped significantly more than expected with known amplified genomic regions, further supporting the non-random nature of their amplification (75 overlaps *versus* 164 non-overlaps for CGAs, 45 overlaps *versus* 194 non-overlaps for random set, p-value = 0.002).

### CGA functional enrichment

A co-expression analysis was performed with gene expression data from all 917 tumor samples, independent of tumor type, using the Hallmark gene set from the Molecular Signature Database (MSigDB)(Liberzon et al. 2015) via the enricher and clusterProfiler R packages. The set of genes comprised 19,768 protein-coding genes and 109 lncRNA-ncCGAs genes. We selected genes expressed in at least 5% of all tumor samples (n samples > 45). A correlation matrix was constructed between lncRNA-CGAs and protein-coding genes, retaining only those Hallmarks that were both strongly and significantly correlated with a given lncRNA (r ≥ 0.15, adjusted p < 0.05). The same analysis was performed for cCGAs for comparison.

### Immunopeptidomics searches

A total of 266 samples from nine publicly available HLA-I immunopeptidomics datasets were analyzed in this study (Tables S20 and S21). The data-dependent acquisition (DDA) files were searched against a personalized, MS-searchable proteome using the FragPipe pipeline (v23.1; https://fragpipe.nesvilab.org/), which incorporates MSFragger (v4.3)(Kong et al. 2017); MSBooster (v1.3.17)(Yang et al. 2023), Percolator (v3.7)(Kall et al. 2007), and Philosopher (v5.1.2)(da Veiga Leprevost et al. 2020).

The search database was constructed by combining the UniProt proteome with isoforms (42,477 entries) and the 235 defined ncCGAs. Default decoy generation methods were applied, and common contaminants were included. The default nonspecific HLA search parameters were modified as follows: DDA+ mode was enabled to improve peptide identification (Yu et al. 2025); only the top 1 peptide per spectrum was reported; ProteinProphet was disabled and Philosopher ion, peptide-spectrum match (PSM), and peptide false discovery rates (FDR) were set to 0.03. Peptide lengths were restricted to 8–15 amino acids.

In addition, we downloaded 494 TimsTOF MS/MS data files from the HLA Ligand Atlas (Marcu et al., 2021), which contain MS/MS measurements of HLA-I binding peptides from various healthy human tissues. We searched these data with Fragpipe using the same parameters and database as above.

### Liquid chromatography and mass spectrometry (LC-MS)

The LC-MS system consisted of an Easy-nLC 1200 coupled to Q Exactive HF-X mass spectrometer (ThermoFisher scientific, Bremen, Germany) and or to Eclipse tribrid mass spectrometer (ThermoFisher Scientific, San Jose, USA). The peptides were eluted on a 450mm analytical column (8 µm m tip, 75 µm m ID) packed with ReproSil-Pur C18 (1.9 µm particle size, 120 A pore size, Dr Maisch, GmbH) and separated at a flow rate of 250 nL/min as described (Chong et al. 2018).

### Parallel reaction monitoring (PRM)

NcCGA-derived peptides with binding affinity (mixMHCpred 3.0; rank z 2%) to any HLA allele of the samples in which they were found (Tadros et al. 2025) or peptides identified with more than 1 peptide spectrum match (PSM) (Table S22) were selected to be validated.

Synthetic peptides were ordered as crude (PTCF, Lausanne, Switzerland). After re-suspension in 2% ACN in 0.1% FA, synthetic peptides were pooled in equimolar concentration of 1pmol/ul and analyzed by PRM. For each peptide charge state of 1+, 2+ and 3+ were monitored (peptide table). For both MS system, the cycle of acquisition consisted of a full survey MS1 scan from 350 – 1650 m/z (R = 60’000, ion accumulation time of 100ms), followed by consecutive acquisition of targed MS/MS scans (R = 30’000, ion accumulation of 60ms).

Collected data was analyzed using Skyline (64-bit, v20.2.0.343), using an ion mass tolerance of 0.05 m/z. Visual inspection of the synthetic and experimental spectra was performed using mirror plots generated with FragPipe-PDV with a tolerance of 20 ppm (Li et al. 2019a). Additionally, three similarity metrics were calculated using the MzJava software library (Horlacher et al. 2015): global cosine score considering all MS/MS peaks in synthetic and endogenous spectra (non-matching peaks set to 0 intensity in complementary spectrum), matched cosine score restricted to matching MS/MS peaks, and the number of matching peaks. Peptides were annotated as “Validated” if their global cosine score was higher than 0.6 or if matched cosine score was higher than 0.6 with more than 15 matched peaks (Table S23). Differences in retention time were also considered and matches were not validated if the difference between the retention time of the endogenous and synthetic peptides was larger than 5 minutes.

One peptide failed synthesis and for five peptides no synthetic spectrum could be found in the PRM files that matched the charge and modification of the endogenous peptides. For these peptides, predicted theoretical spectra calculated by MSBooster, using DIA-NN deep learning models, from FragPipe and visualized with FragPipe-PDV, in addition to hyperscore, were used to compare them (Table S24).

### Statistical tests and figures

Statistical analyses were performed using R (v4.1.2). Comparisons between medians were performed using Wilcoxon test (Fig. 1D). The difference between two proportions was assessed using Fisher’s exact tests. The p value adjustment of the enrichment analysis were performed with the Bonferroni method (Fig. 5). Correlations are calculated with the Pearson method.

### Data and materials availability

All data supporting the findings of this study are provided within the paper and its Supplementary Materials. Supplementary Tables have been deposited in the Figshare repository: https://doi.org/10.6084/m9.figshare.32310897

## Acknowledgements

We acknowledge cell drawing from Figure 1A by Léa Lortal from the Noun Project and organs by Adrián Juárez Granados. We are grateful to Pablo Sarobe for discussions on this work. This work was funded by Ministerio de Ciencia, Innovación y Universidades grant PID2021-128791OB-I00, PID2024-162742OB-I00, PID2021-122726NB-I00 and PRE2022-101505 fellowship for N.K., funded by MCIN/AEI/10.13039/501100011033 and by “ERDF: A way of making Europe”, by the “European Union”. We also acknowledge funding from Generalitat de Catalunya, grant 2021SGR00042, Gobierno de Navarra, grant 0011-1411-2024-000077 and the Instituto de Salud Carlos III, which finances Centro de Investigación Biomédica en Red de Enfermedades Hepáticas y Digestivas (CIBEREhd), financed by the EU (NextGenerationEU), Plan de Recuperación Transformación y Resiliencia). The work was also funded by the European Union (ERC, NovoGenePop, project number 101052538). Views and opinions expressed are however those of the authors only and do not necessarily reflect those of the European Union or the European Research Council. Neither the European Union nor the granting authority can be held responsible for them. The work was further funded by the Ludwig Institute for Cancer Research.

## Author contributions

M.M.A., J.P-B., and M.E.C conceptualized the project and interpreted the results. M.E.C compiled and processed the samples and performed most computational analyses. J.C.M. set up and automated the ribosome profiling computational framework. M.E.C., C.V., and C.P. performed the evolutionary analyses. S.R.-S. contributed with computational analysis of the RiboSeq data. N.K. and J.C.G.-S contributed data analyses. M.T.-C. and H.P. conducted the processing and measurement of PRM LC-MS experiments. M.M. and M.E.C performed MS search analysis and MS/MS comparisons. M.M.A., J.P-B., P.F. and M.B-S. supervised the study. M.M.A. and M.E.C wrote the manuscript and generated the figures with contributions from all authors. All authors contributed to the interpretation of the results and commented on the manuscript.

