## Supplementary Figures for "Identification of a large class of cancer–germline microproteins as a source of immunotherapeutic targets"

**Supplementary Figures**
Camarena et al.

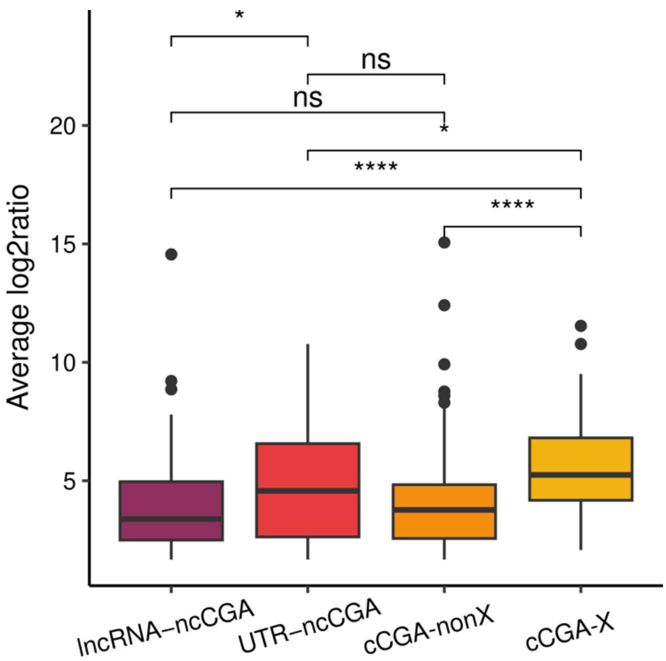


**Figure S1. CGA expression in tumor *versus* expression in adjacent normal tissue.** Shown is the distribution of $log_2$ of the ratio of the expression (in transcripts per million or TPM) in tumor versus normal tissue for each patient and transcript encoding a CGA, classified by CGA type. Data is for 917 patients with matched tumor/normal RNA-Seq data. Selected genes contained were exclusively translated in testis, not expressed in GTEX somatic tissues, and showed an enrichment in tumor *versus* adjacent normal tissue (tumor > 1 TPM and $log_2$ TPM tumor/ TPM adjacent > 1.585 in at least one patient sample). X refers to genes located in chromosome X, nonX to genes located in autosomes.

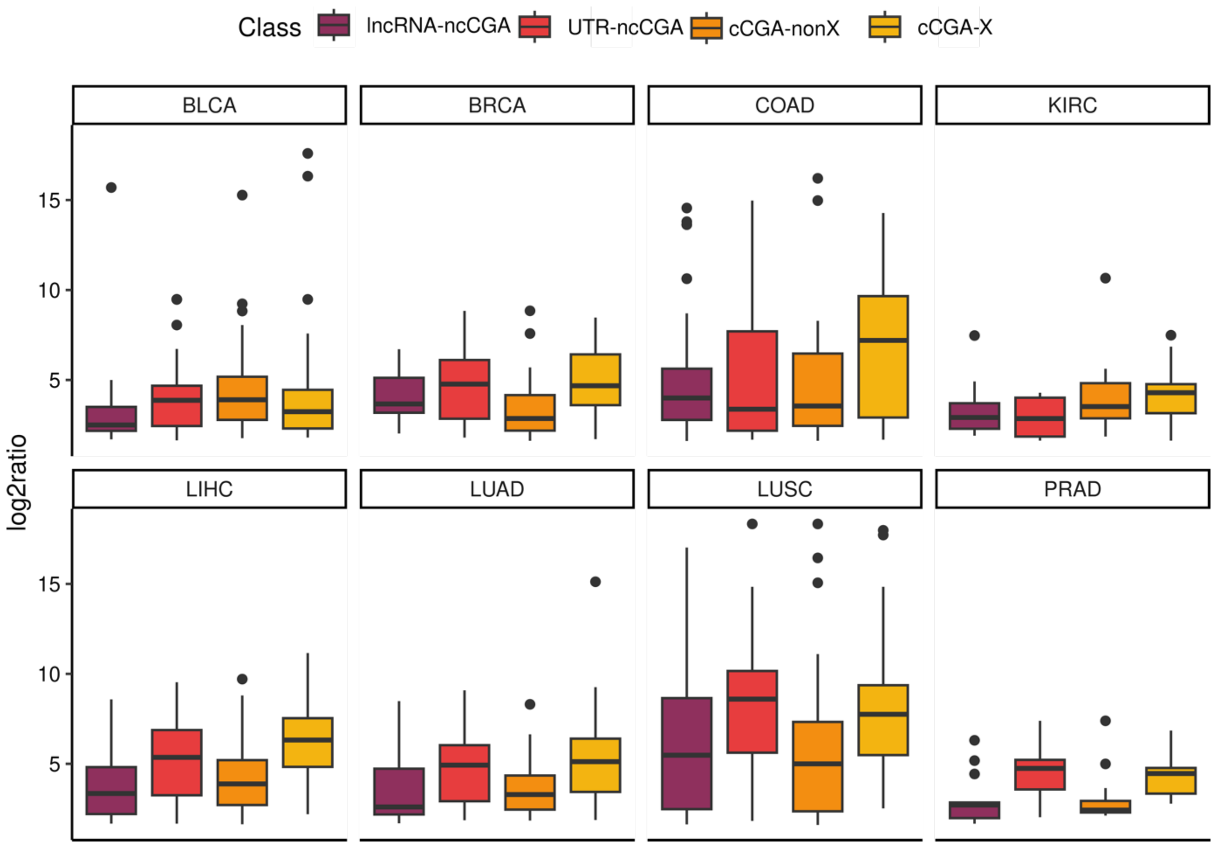


**Figure S2. Expression in tumor *versus* expression in adjacent normal tissue for CGA transcripts per cancer type.** Same data as in Figure S1 but separated by cancer type.

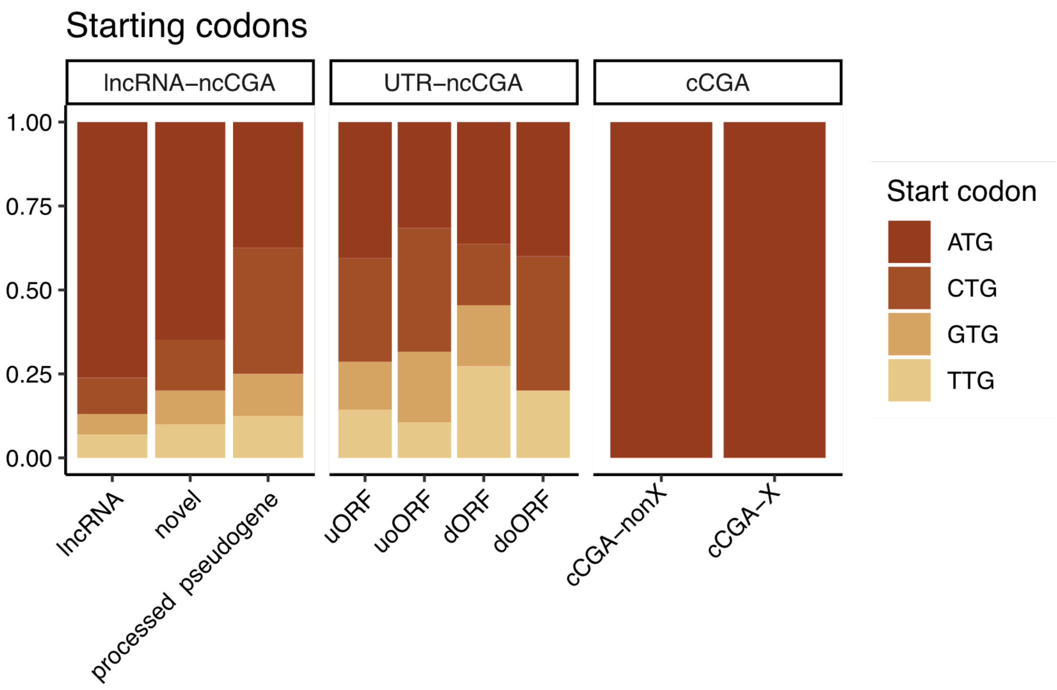


**Figure S3. Start codon distribution for different CGAs types.** Number of CGAs of different types: X cCGA 66, nonX cCGA 128, lncRNA-ncCGA lncRNA 130, lncRNA-ncCGA novel 20, lncRNA-ncCGA processed pseudogene 8, UTR-ncCGA uORFs 42, UTR-ncCGA uoORFs 19, UTR-ncCGA dORFs 11, UTR-ncCGA doORFs 5.

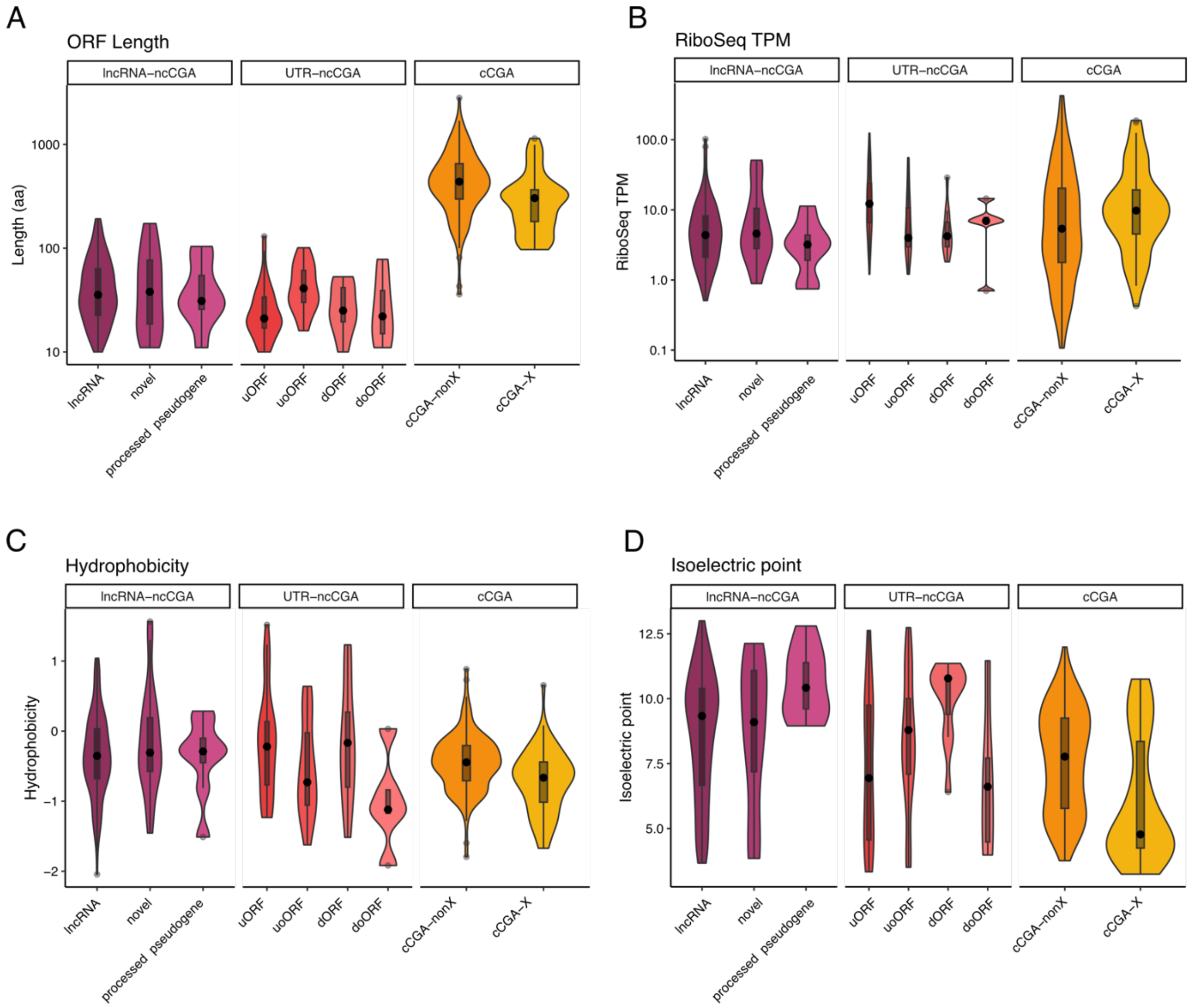


**Figure S4. Characteristics of CGAs by subtype.** Distribution of amino acid sequence length, translation level (Ribo-Seq TPM), isoelectric point and hydrophobicity for different classes of translated ORFs.

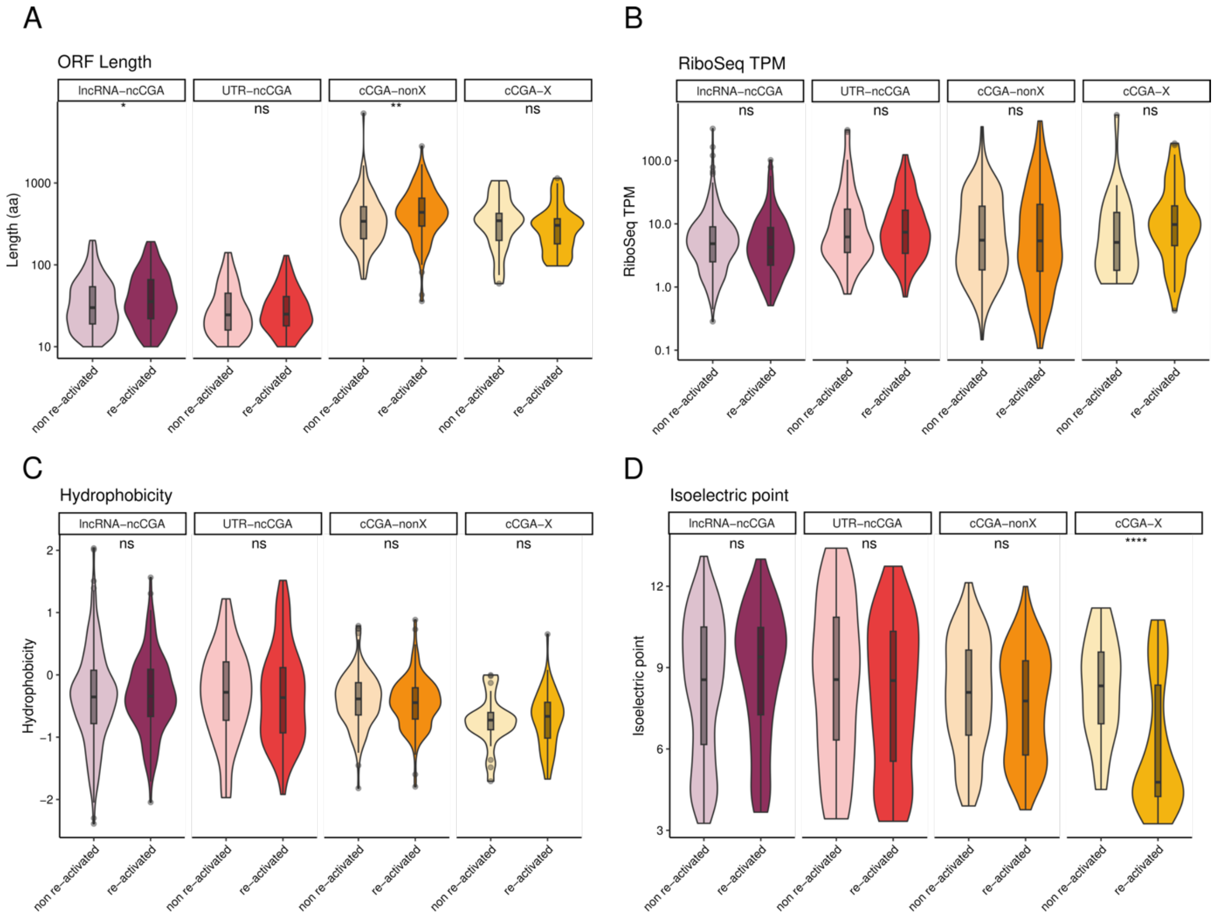


**Figure S5. Features of testis translated ORFs depending on whether they are reactivated in tumors or not.** Non re-activated refers to the set of translated ORFs that is testis-specific but not detected in tumor samples. Reactivated refers to the set not expressed in GTEX somatic tissues, and showing an enrichment in tumor *versus* adjacent normal tissue (tumor > 1 TPM and $\log_2$ TPM tumor/ TPM adjacent > 1.585 in at least one patient sample).

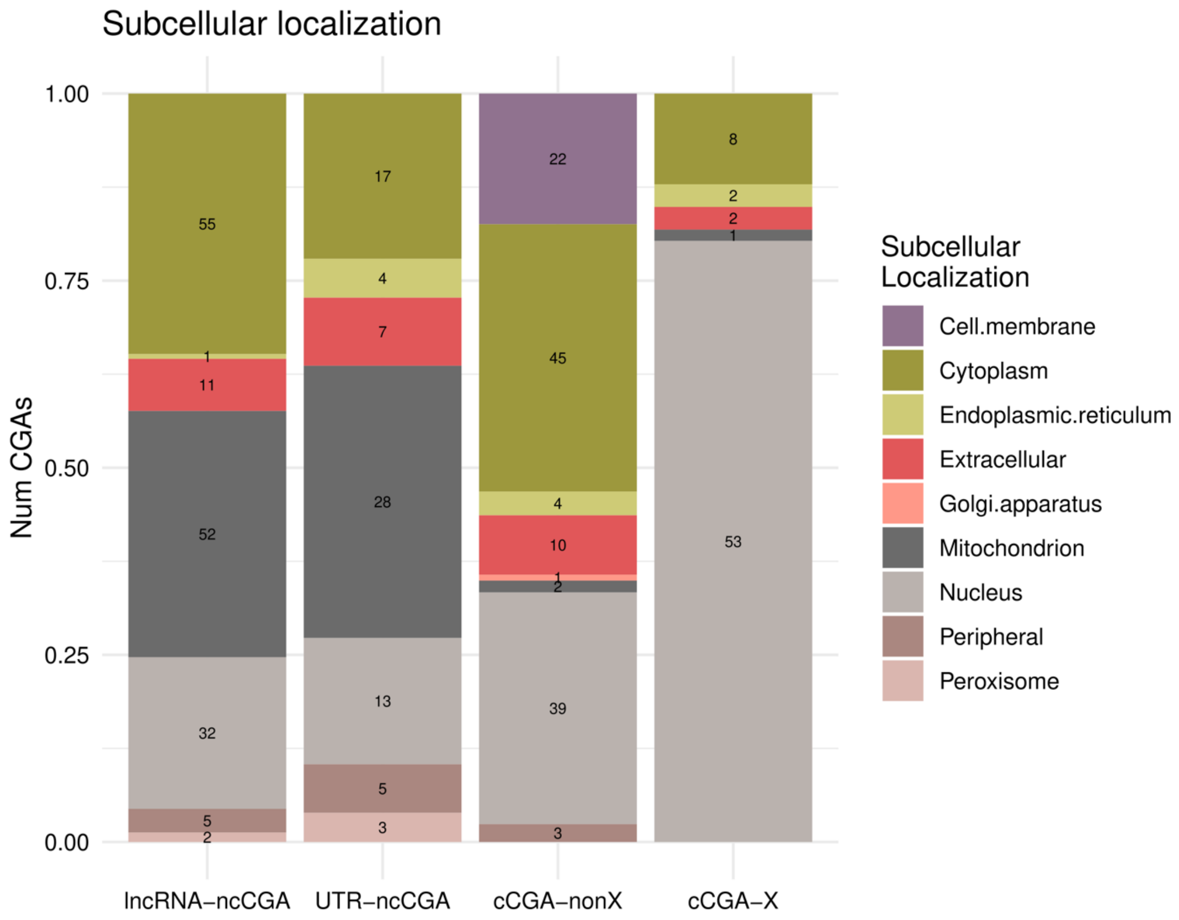


**Figure S6. Subcellular location predictions for CGAs.** Predictions were performed with the DeepLoc 2.1 software. If several localizations were reported, we chose the one with the highest probability assigned

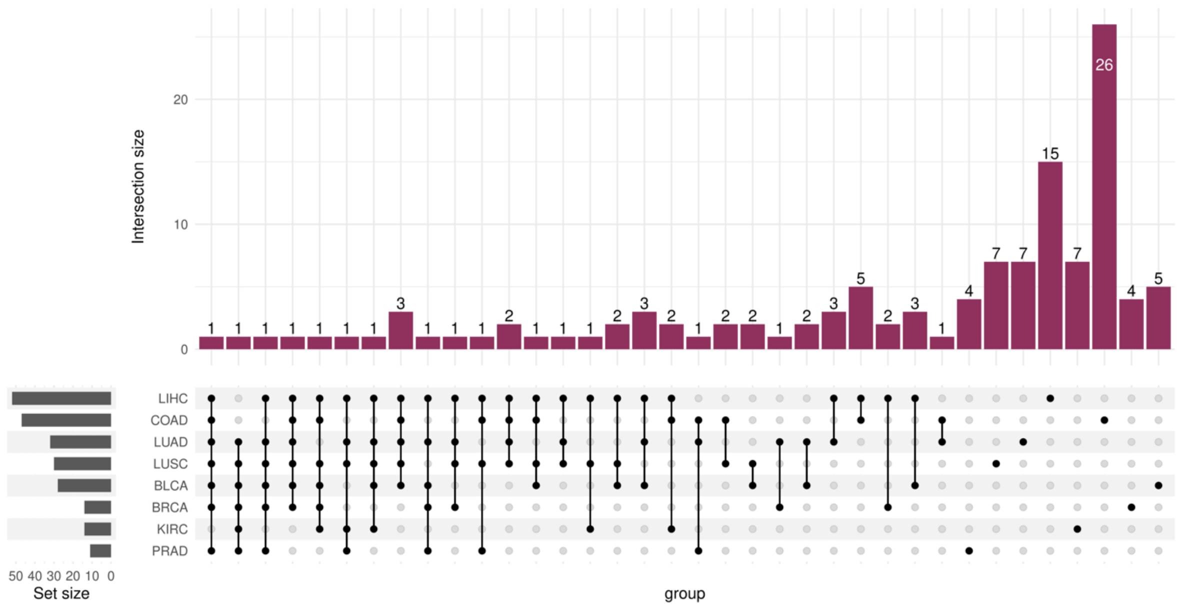


**Figure S7. Distribution of lncRNA-ncCGAs across tumor types.** Data is shown at the level of the genes. 47 lncRNAs containing translated ORFs are transcribed in more than one cancer type, while 75 are cancer-type specific. COAD is the cancer type with the highest number of lncRNA-ncCGAs, but in general they are patient-specific (supplementary data).

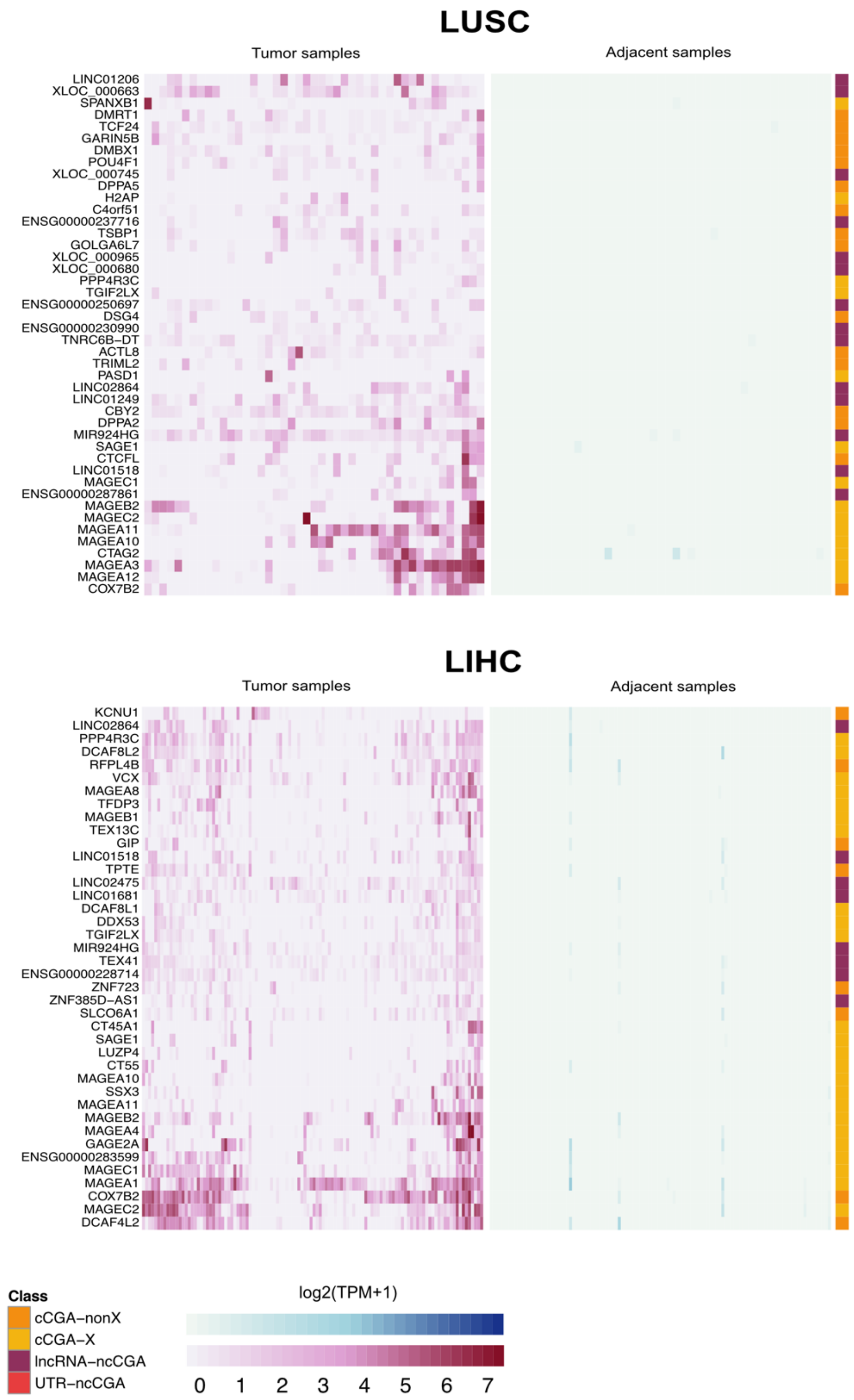


**Figure S8. Distribution of CGAs across patients in tumor and adjacent samples in LUSC and LIHC.** The CGAs shown have been found in more than 10% of the tumor samples per cancer type and display high tumor-specificity. In particular, they are expressed in more than 10% of the tumor samples at TPM > 1 and in adjacent < 0.1 TPM, and this level of specificity is fulfilled by at least 90% of the patients. LUSC: lung squamous cell carcinoma; LIHC: liver hepatocellular carcinoma.

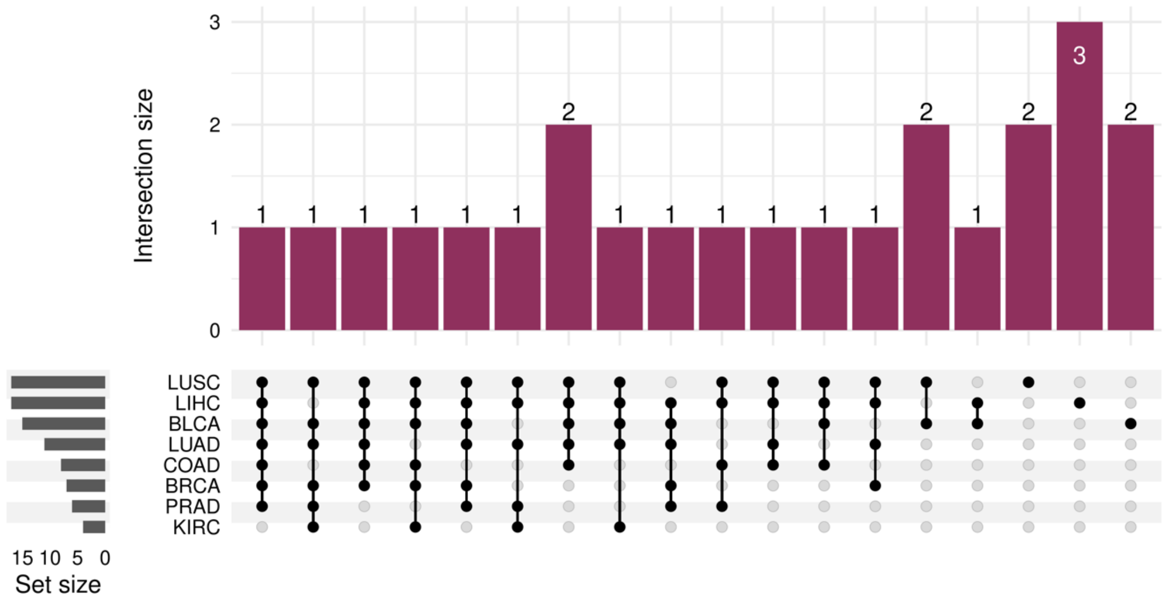


**Figure S9. Distribution of lncRNA-ncCGAs across tumor types for genes found in > 10% of the patients in at least once cancer type.** As in Figure S7 but for the set of most widely found lncRNA-ncCGAs.

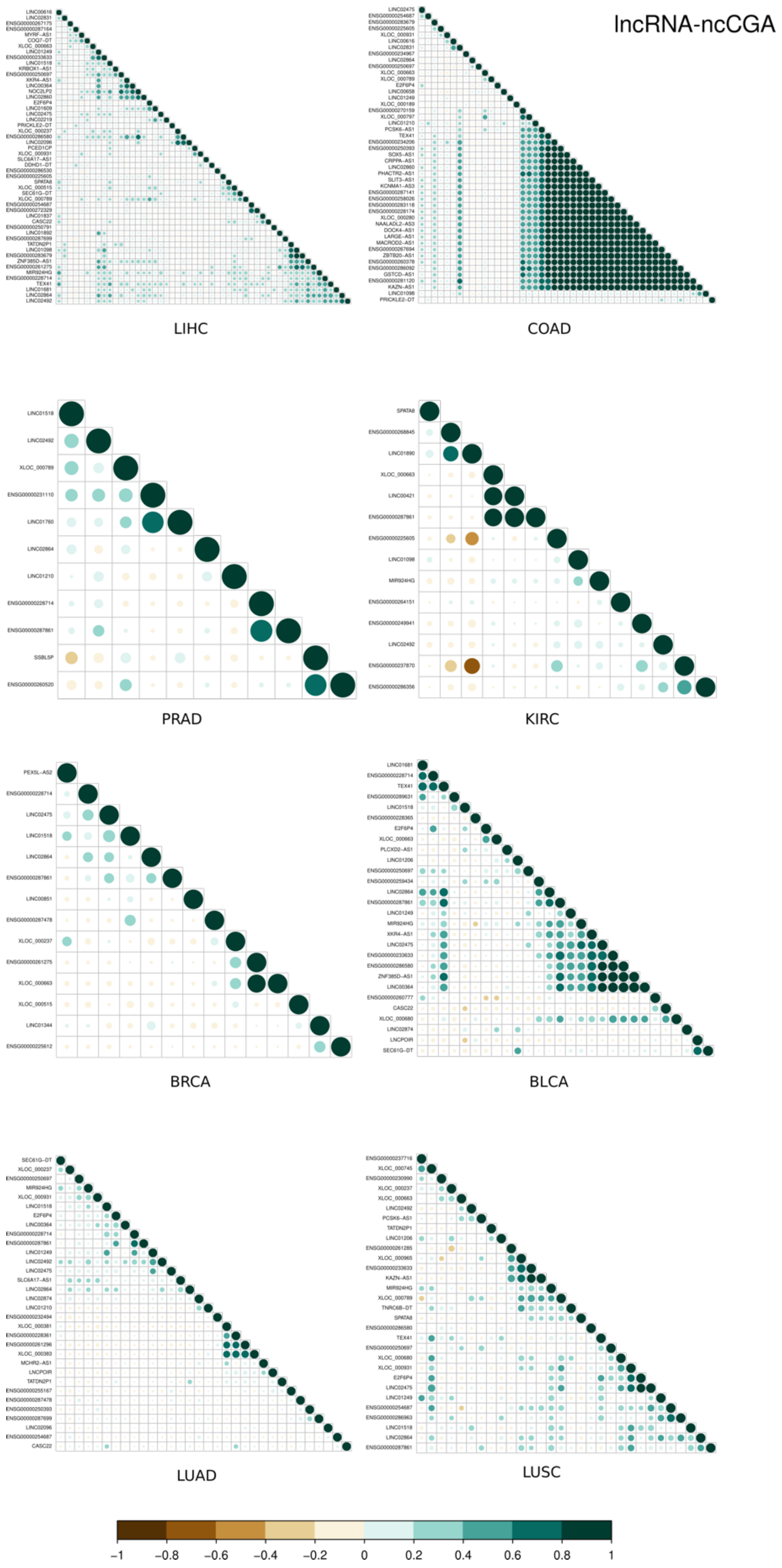


**Figure S10. Expression correlation of lncRNA-ncCGA gene expression.** Correlation values are shown in a scale from -1 to 1. In most cancer types the expression correlation values are low, denoting that different patients tend to express a different subset of lncRNA-ncCGAs.

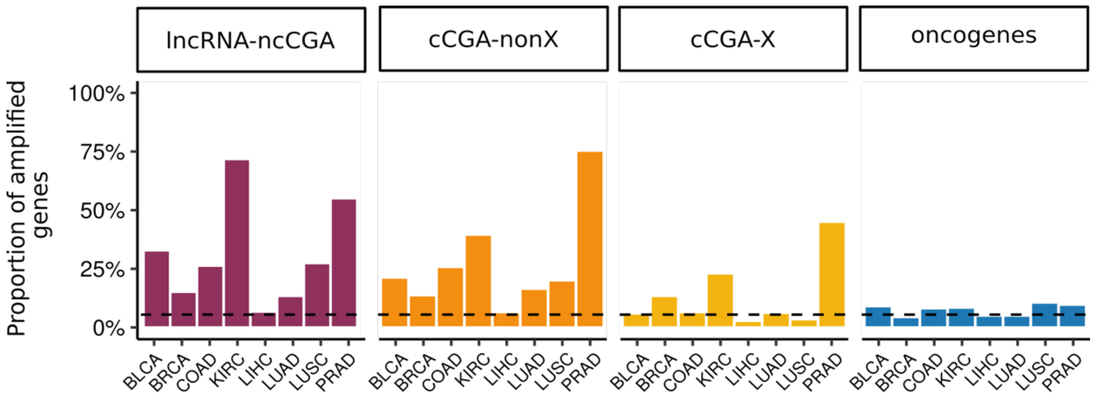


**Figure S11. Percentage of CGAs/oncogenes amplified in different cancer types.** As in Figure 4B but in %.

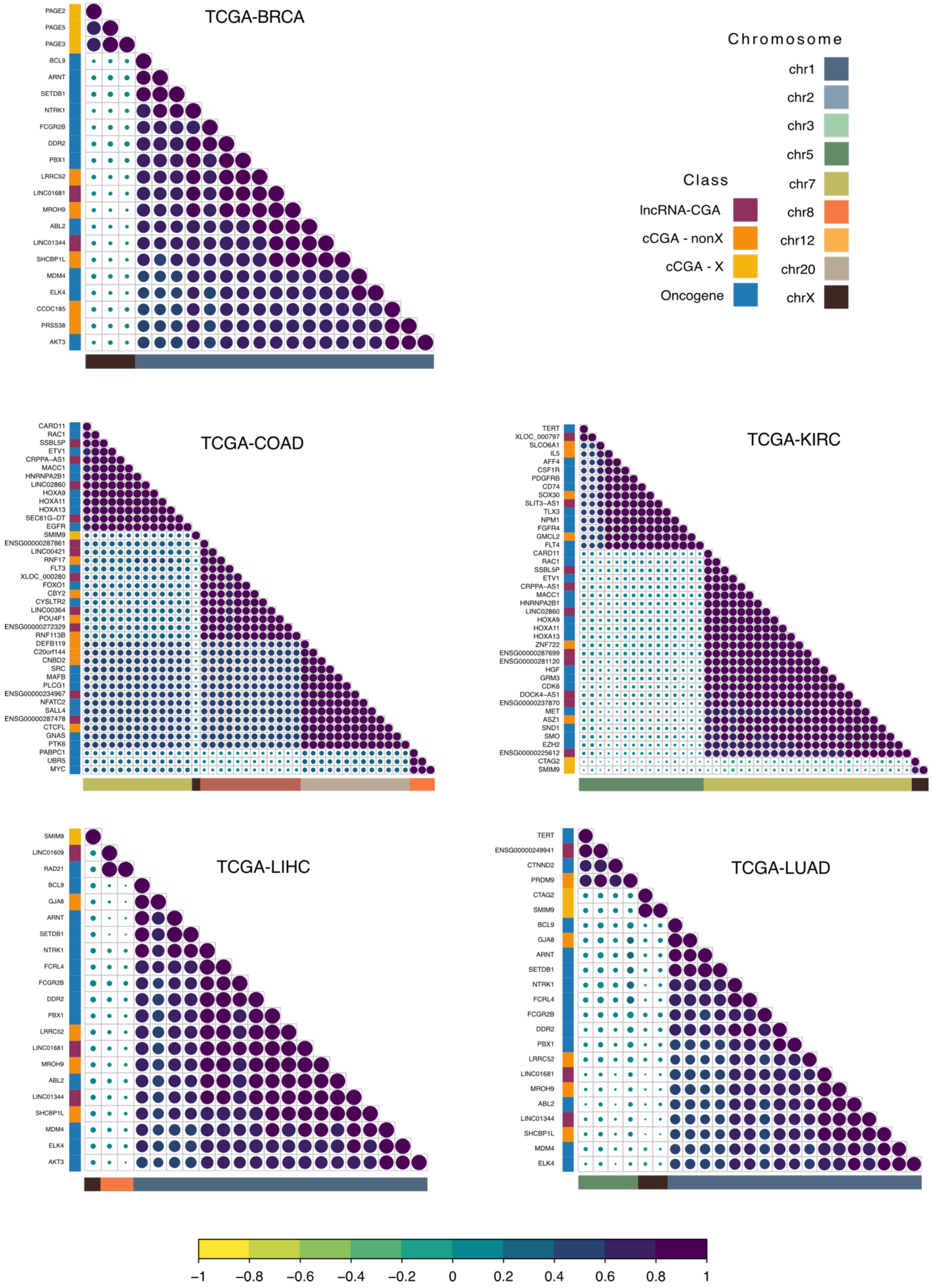


**Figure S12. Blocks of co-amplified genes in different cancer types**. Plots are as in Figure 4C but for other cancer types.

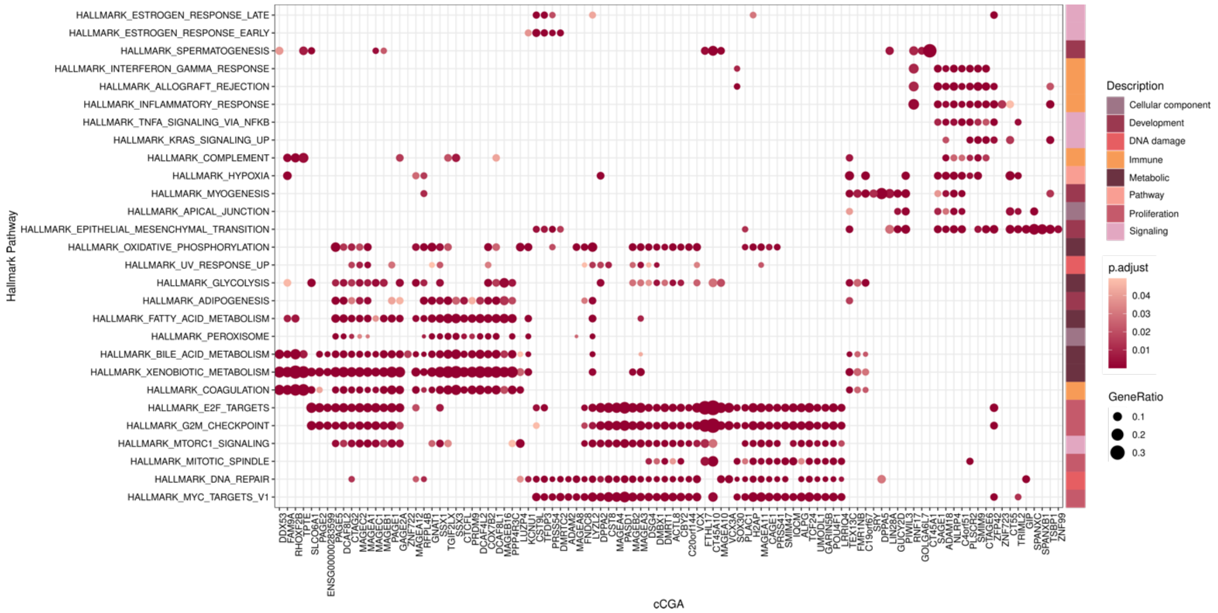


**Figure S13. Hallmark gene set associations based on co-expression for cCGAs**. Similar to Figure 5 but for cCGAs

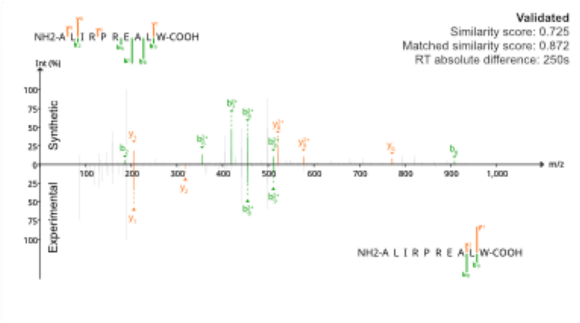
Validated
Similarity score: 0.725
Matched similarity score: 0.872
RT absolute difference: 250s
NH2-A L I R P R E A L W-COOH
Int (%)
Synthetic
Experimental
m/z

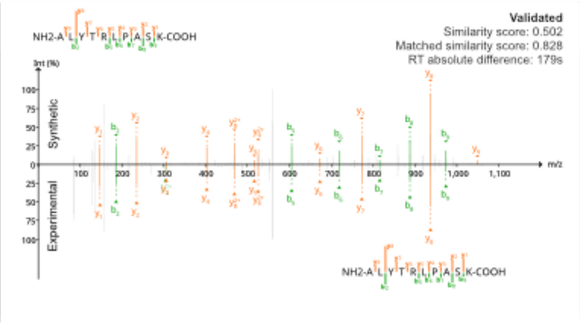
Validated
Similarity score: 0.502
Matched similarity score: 0.828
RT absolute difference: 179s
NH2-A L Y T R L P A S K-COOH
Int (%)
Synthetic
Experimental
m/z

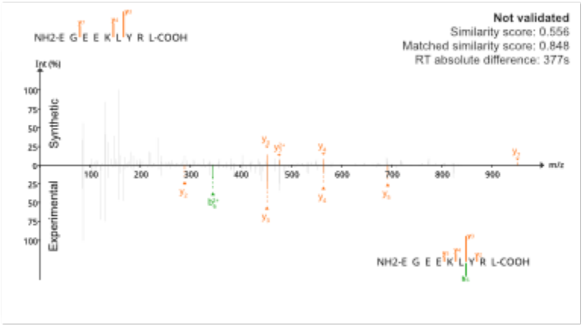
Not validated
Similarity score: 0.556
Matched similarity score: 0.848
RT absolute difference: 377s
NH2-E G E E K L Y R L-COOH
Int (%)
Synthetic
Experimental
m/z

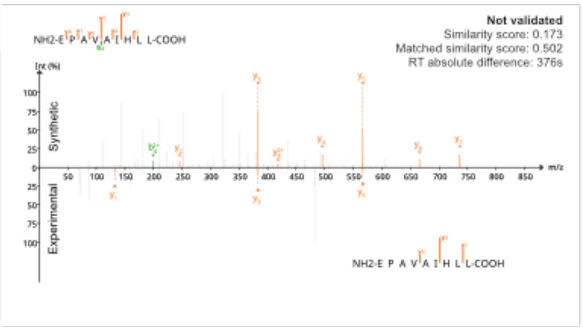
Not validated
Similarity score: 0.173
Matched similarity score: 0.502
RT absolute difference: 376s
NH2-E P A V A I H L L-COOH
Int (%)
Synthetic
Experimental
m/z

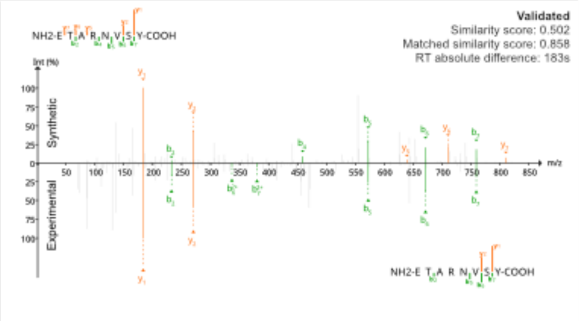
Validated
Similarity score: 0.502
Matched similarity score: 0.858
RT absolute difference: 183s
NH2-E T A R N V S Y-COOH
Int (%)
Synthetic
Experimental
m/z

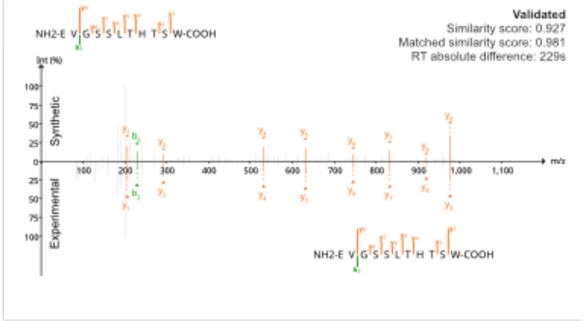
Validated
Similarity score: 0.927
Matched similarity score: 0.981
RT absolute difference: 229s
NH2-E V G S S L T H T S W-COOH
Int (%)
Synthetic
Experimental
m/z

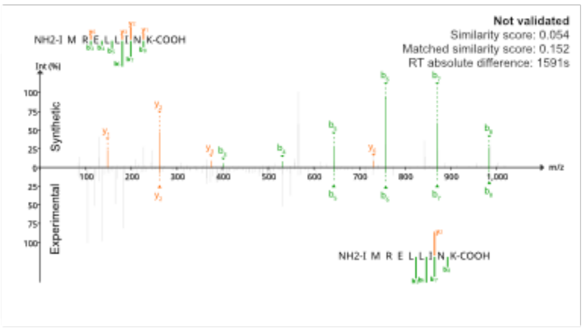
Not validated
Similarity score: 0.054
Matched similarity score: 0.152
RT absolute difference: 1591s
NH2-I M R E L L I N K-COOH
Int (%)
Synthetic
Experimental
m/z

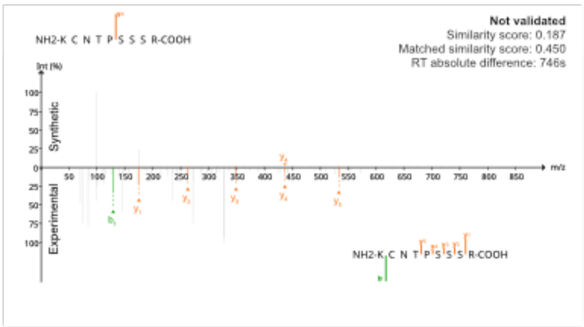
Not validated
Similarity score: 0.187
Matched similarity score: 0.450
RT absolute difference: 746s
NH2-K C N T P S S S R-COOH
Int (%)
Synthetic
Experimental
m/z

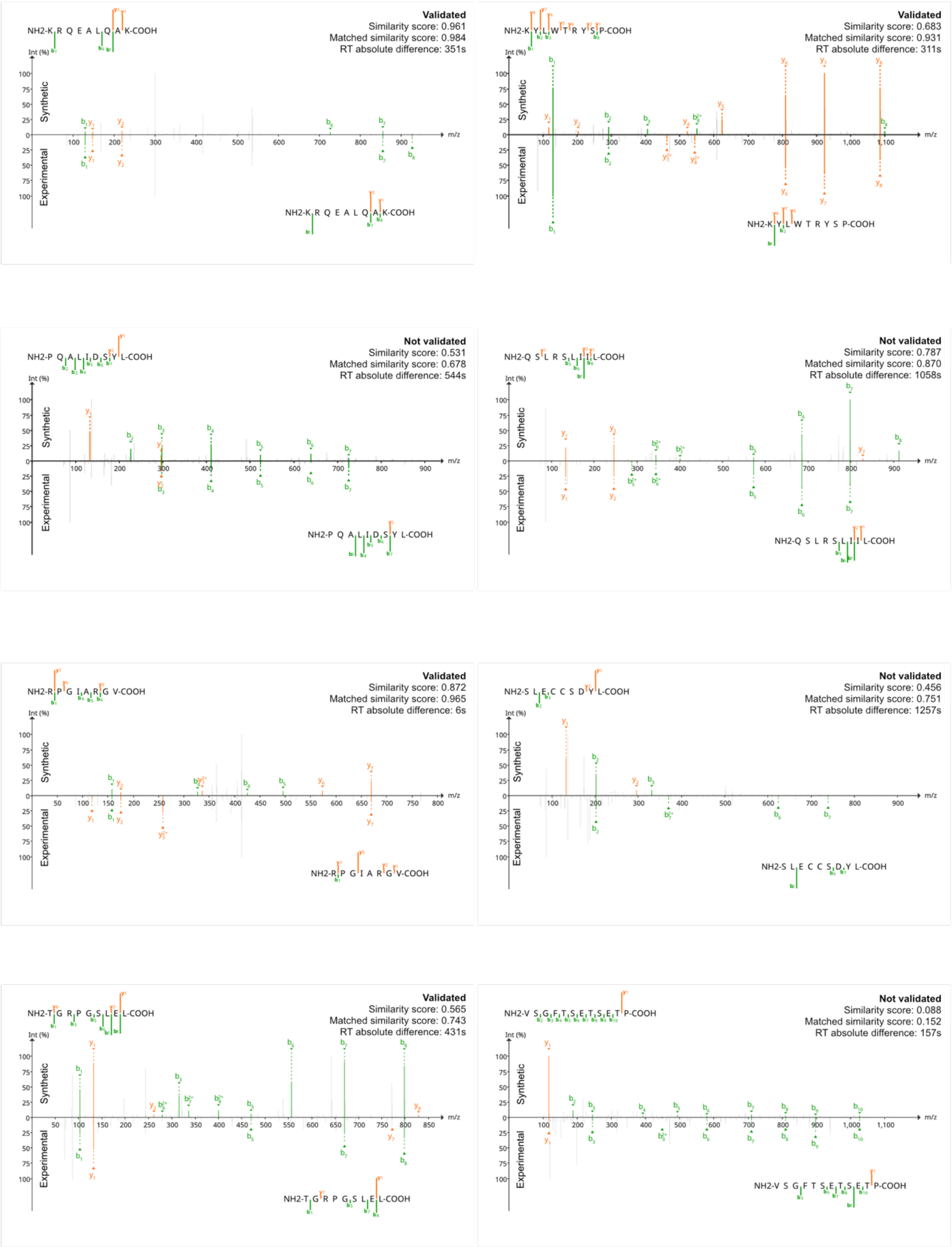
NH2-K R Q E A L Q A K-COOH
Validated
Similarity score: 0.961
Matched similarity score: 0.984
RT absolute difference: 351s
NH2-K Y L W T R Y S P-COOH
Validated
Similarity score: 0.683
Matched similarity score: 0.931
RT absolute difference: 311s
NH2-P Q A L I D S Y L-COOH
Not validated
Similarity score: 0.531
Matched similarity score: 0.678
RT absolute difference: 544s
NH2-Q S L R S L I I L-COOH
Not validated
Similarity score: 0.787
Matched similarity score: 0.870
RT absolute difference: 1058s
NH2-R P G I A R G V-COOH
Validated
Similarity score: 0.872
Matched similarity score: 0.965
RT absolute difference: 6s
NH2-S L E C C S D Y L-COOH
Not validated
Similarity score: 0.456
Matched similarity score: 0.751
RT absolute difference: 1257s
NH2-T G R P G S L E L-COOH
Validated
Similarity score: 0.565
Matched similarity score: 0.743
RT absolute difference: 431s
NH2-V S G F T S E T S E T P-COOH
Not validated
Similarity score: 0.088
Matched similarity score: 0.152
RT absolute difference: 157s
Int (%)
Synthetic
Experimental
m/z

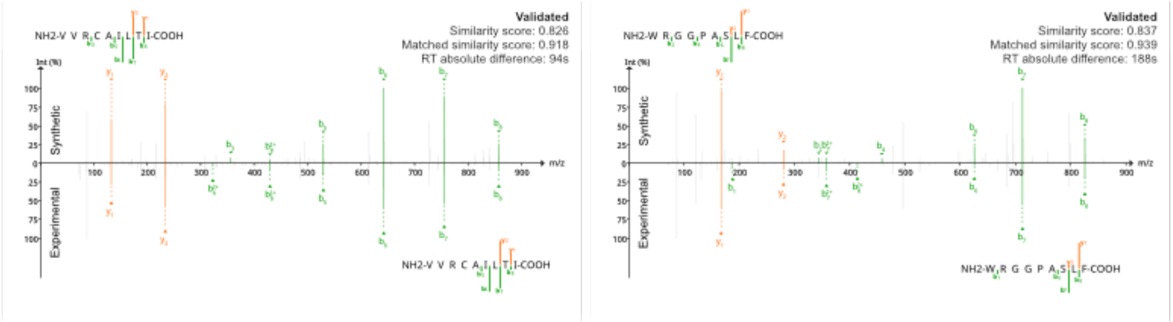


**Figure S14. Validation of non-canonical peptides with synthetic peptides**.
The above MS spectra correspond to the spectra of the synthetic peptide while the below are the MS spectra of the experimental peptide. Additional information is provided in the supplementary table 23.

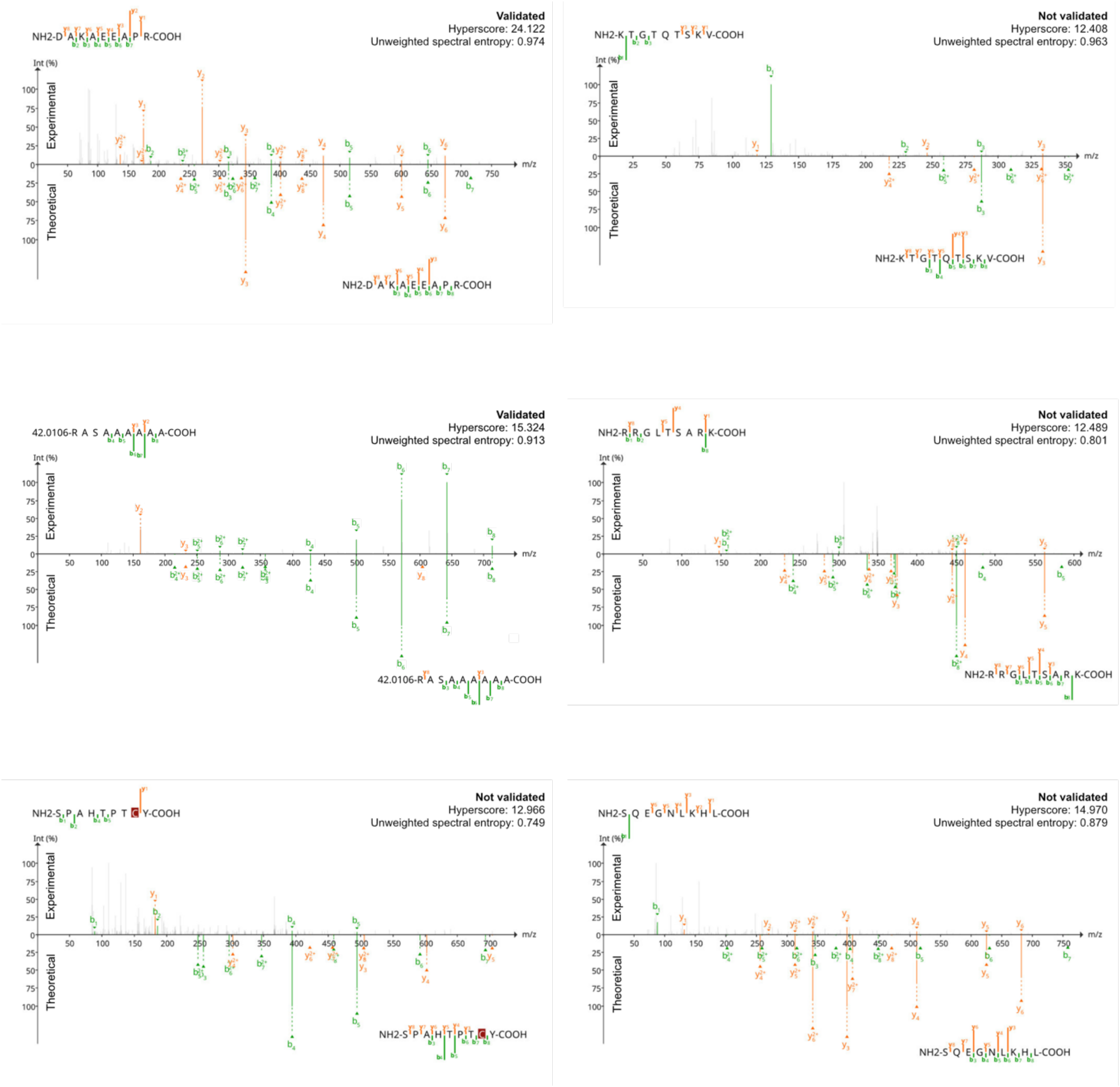


**Figure S15. Validation of non-canonical peptides with predicted theoretical spectra**. The above MS spectra correspond to the spectra of the experimental peptide while the below are the theoretical MS spectra. Additional information is provided in the supplementary table 24.

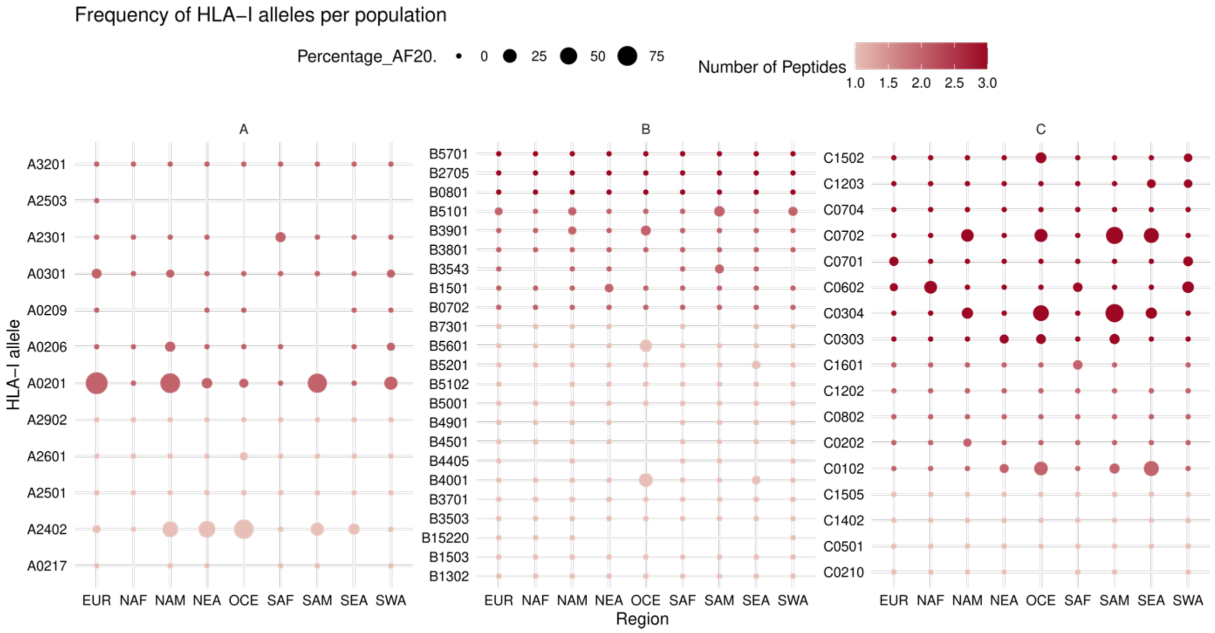


**Figure S16. Population frequencies of HLA-I alleles predicted to present validated ncCGA-derived peptides with immunogenic potential according to PRIME 2.1.** The color gradient refers to the number of validated peptides whereas the size of the point refers to the proportion of populations in which that specific allele has an allele frequency (AF) > 20.

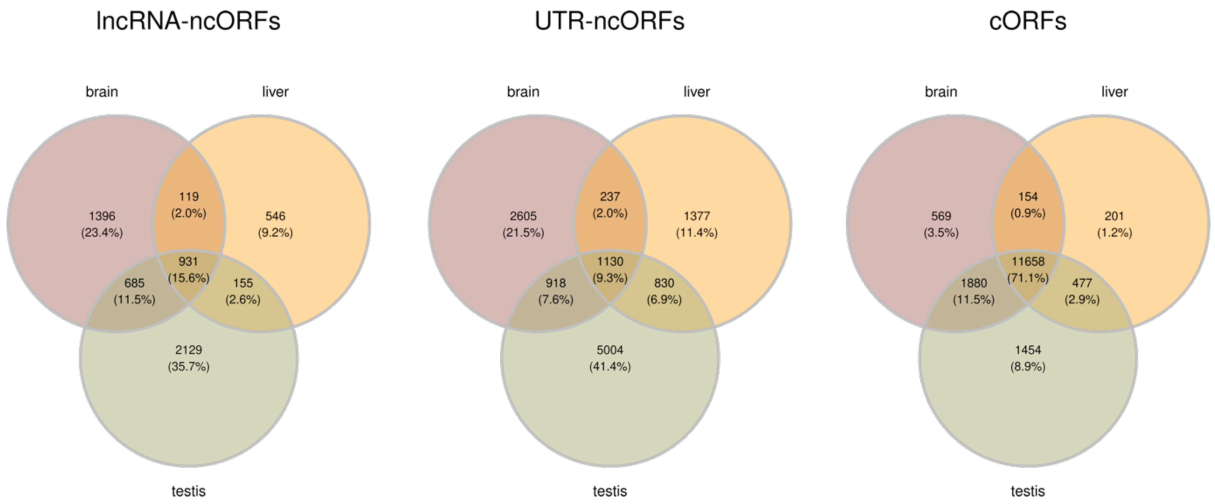


**Figure S17. Analyses of translated ORFs in different tissues.** Shown is the number of ORFs detected in different tissues and their intersections. Translation in a sample from a given tissue was considered a positive for that tissue. cORFs: canonical ORFs (annotated coding sequences). ncORFs: non-canonical ORFs.

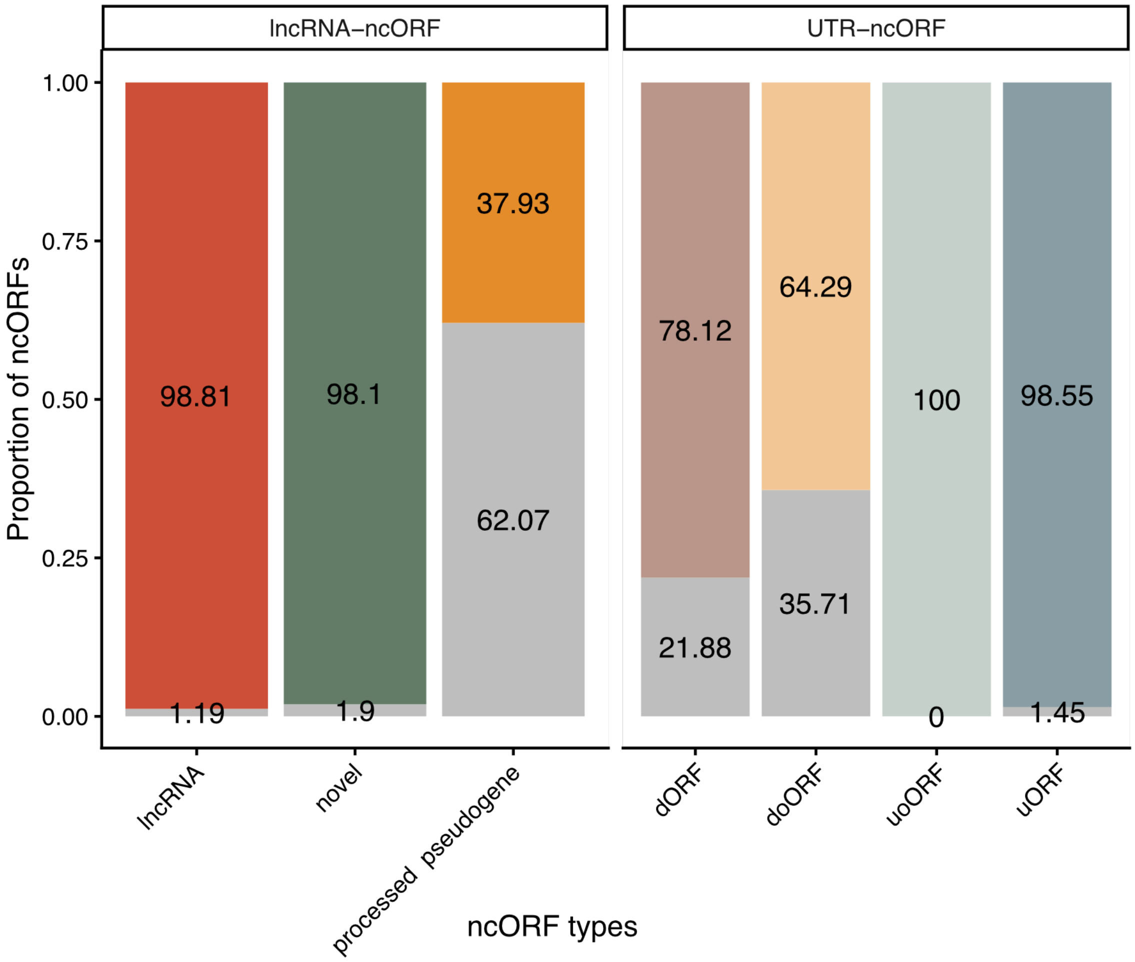


**Figure S18. Proportion of translated ncORFs with homology to annotated human proteins.** In grey the percentage of protein products from ncORFs that had homologues in the human genome using BLASTP searches and an E-value < $10^{-4}$.

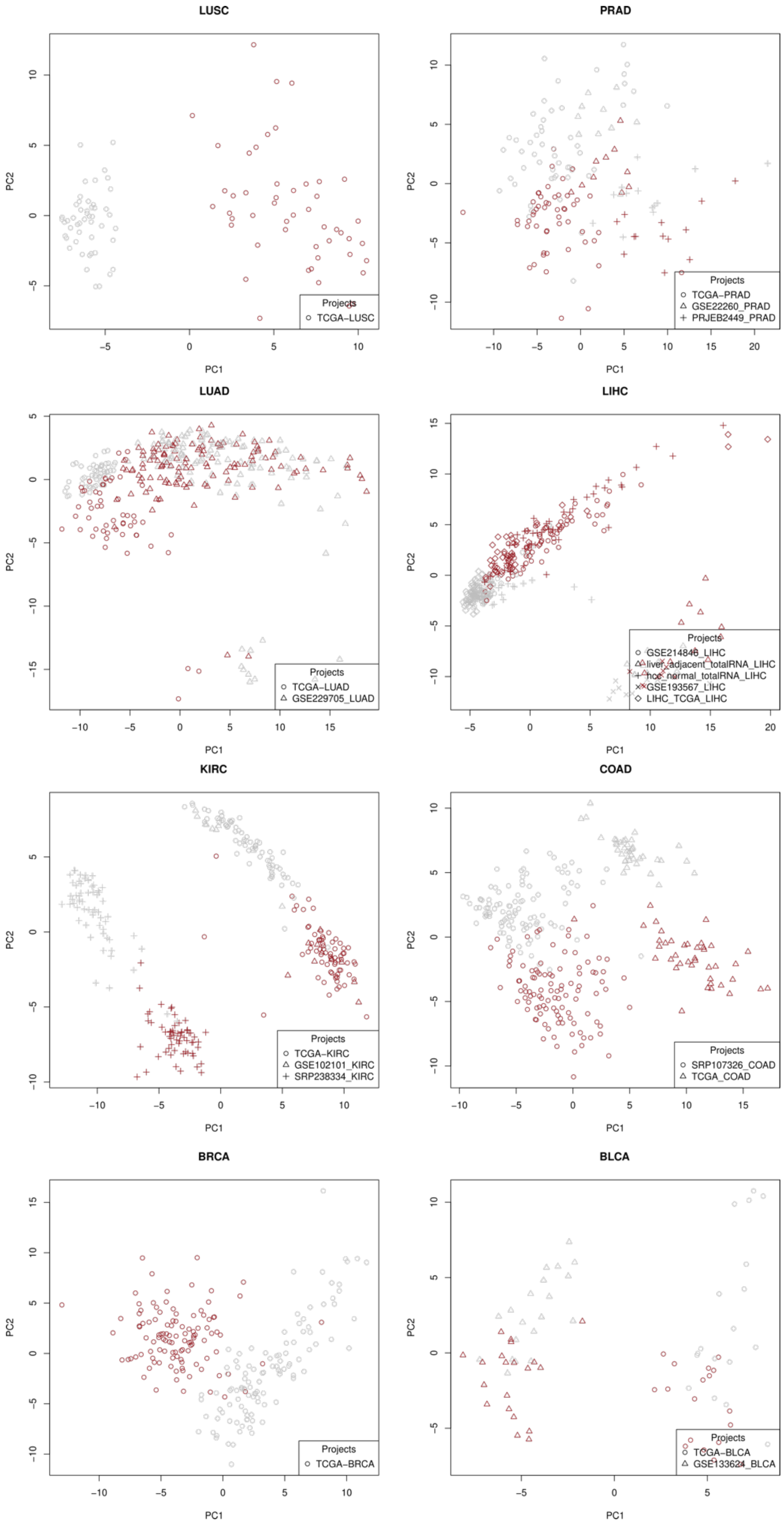


**Figure S19. Principal component analysis (PCA) of gene expression data for different cancer types**. The PCA was performed with the 100 most variable genes per cancer type to validate that adjacent and tumor samples were properly separated.

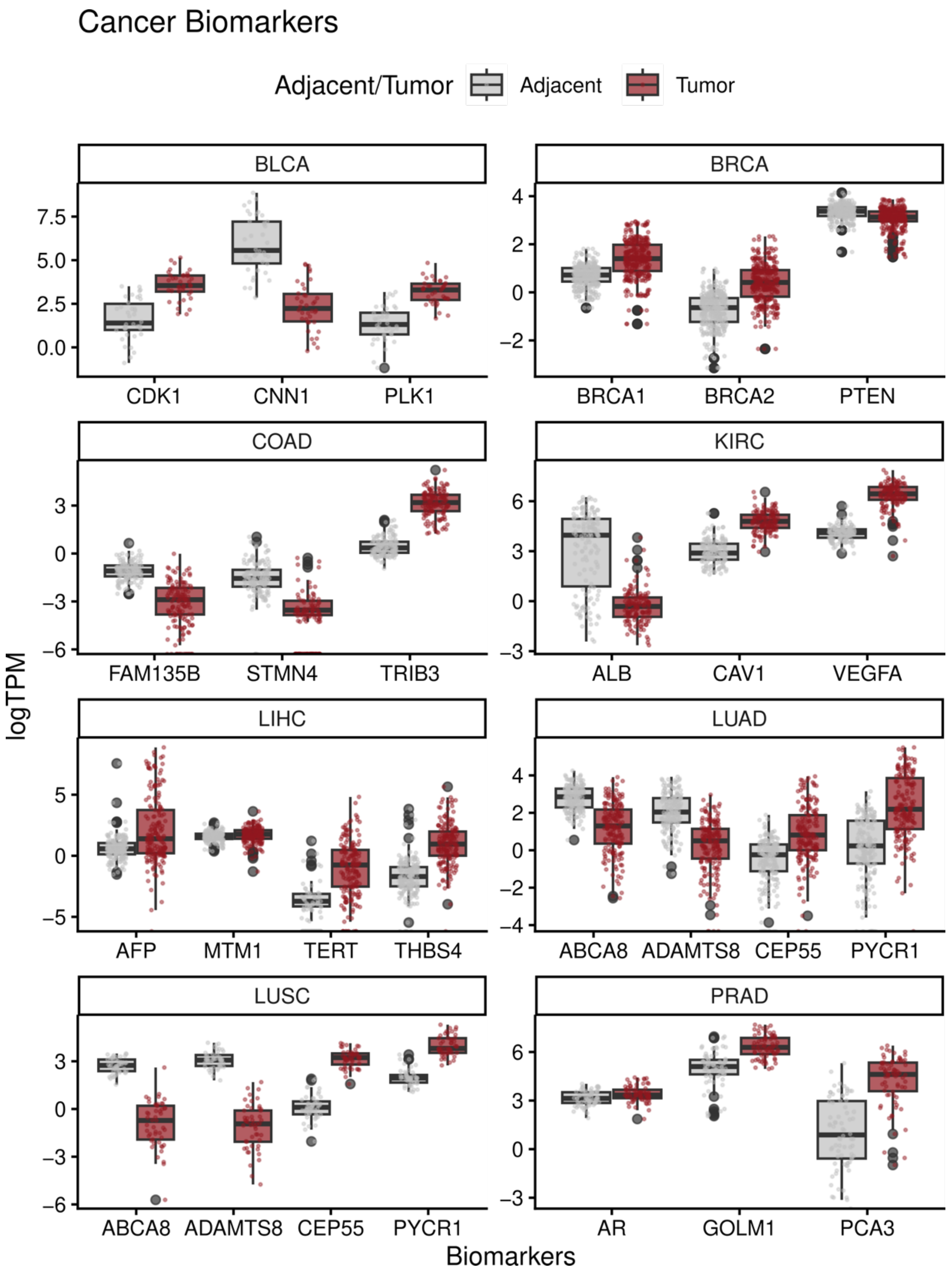


**Figure S20. Gene expression values for cancer biomarkers in tumor and normal adjacent samples.** The data shows the expected biases in the distribution of these markers in the two types of samples.
